# Human Brain-Wide Activation of Sleep Rhythms

**DOI:** 10.1101/2024.10.14.618165

**Authors:** Haiteng Wang, Qihong Zou, Jinbo Zhang, Jia-Hong Gao, Yunzhe Liu

## Abstract

During sleep, our brain undergoes highly synchronized activity, orchestrated by distinct neural rhythms. Little is known about the associated brain activation during these sleep rhythms, and even less about their functional implications. In this study, we investigated the brain-wide activation underlying human sleep rhythms by employing simultaneous electroencephalography (EEG) and functional magnetic resonance imaging (fMRI) in 107 participants during nocturnal nap (first half of the night). We identified robust coupling between slow oscillations (SOs) and fast spindles during deep non-rapid eye movement (NREM) sleep (N2/3 stages), with spindle peaks consistently occurring just before the SO UP-state. This SO-spindle coupling was linked to elevated activation in both the thalamus and hippocampus, alongside increased functional connectivity from the hippocampus to the thalamus and from the thalamus to the medial prefrontal cortex (mPFC). An open-ended cognitive state decoding analysis suggested that these activations may relate to episodic memory processes, yet were distinct from task-related networks. Together, these findings highlight the thalamus as a key coordinator of hippocampal-cortical communication during sleep and provide new insights into the mechanisms by which synchronized sleep rhythms may support memory consolidation.

## Introduction

During sleep, the human brain engages in highly synchronized activity driven by distinct neural rhythms. In non-rapid eye movement (NREM) sleep, the emergence and coordination of slow oscillations (SOs) and sleep spindles are key features (Hahn et al., 2020; Helfrich et al., 2018; Ngo, Fell, & Staresina, 2020; Schreiner et al., 2022; Staresina et al., 2015; Staresina et al., 2023). However, the brain activation patterns associated with these rhythms, and especially their coordination, remain largely unexplored, and their functional implications are even less understood.

SOs (∼1 Hz) are characterised by fluctuations in neuronal membrane potential between periods of silence (’hyperpolarization’, or ‘DOWN’ state) and excitation (’depolarization’, or ‘UP’ state) (Amzica & Steriade, 2002; Steriade, McCormick, & Sejnowski, 1993). These oscillations originate in neocortical regions, particularly the medial prefrontal cortex (mPFC), during NREM sleep (Dang-Vu et al., 2008; Massimini et al., 2004; Nir et al., 2011). The depolarization phase of SOs is believed to trigger sleep spindles in the thalamus, producing 11–16 Hz oscillations (Fernandez & Lüthi, 2020; Mak-McCully et al., 2017). Thalamic spindles tend to align with the excitable UP-states of SOs and can synchronize with hippocampal ripples (Buzsáki, 2015; Helfrich et al., 2019; Joo & Frank, 2018; Ngo, Fell, & Staresina, 2020). Rodent optogenetics studies have shown that only thalamic spindles phase-locked to the UP-state of cortical SOs enhance memory consolidation, while out-of-phase spindles do not (Latchoumane et al., 2017). This finding is consistent with intracranial evidence from human epilepsy patients, suggesting the thalamus acts as a key relay between the hippocampus and cortex, coordinating their interaction (Coulon, Budde, & Pape, 2012; Ferraris et al., 2021; Schreiner et al., 2022; Staresina et al., 2015).

The triple coupling of SOs, spindles, and ripples is believed to facilitate memory consolidation by synchronizing neuronal activity across brain regions and replaying memory traces from the hippocampus to the neocortex (Diekelmann & Born, 2010; Latchoumane et al., 2017; Liu et al., 2022; Singh, Norman, & Schapiro, 2022; Staresina et al., 2015; Staresina et al., 2023), possibly through relay from thalamic spindles. Consistent with this view, prior human EEG-fMRI studies have reported spindle-related activation not only in the thalamus, but also in the hippocampus and adjacent parahippocampal gyrus (Bergmann et al., 2012; Schabus et al., 2007). Spindle-related activity has also been linked to striatal engagement, suggesting a broader network that may support memory-related processing during sleep (Fogel et al., 2017). However, the exact mechanisms of this inter-regional communication during coordinated sleep rhythms remain unclear. Understanding the brain-wide activation associated with these rhythms is crucial, as it reveals how information flows among brain regions, especially during the critical time windows when these rhythms align.

Investigating these sleep-related neural processes in humans is challenging because it requires tracking transient sleep rhythms while simultaneously assessing their widespread brain activation. Recent advances in simultaneous EEG-fMRI techniques provide a unique opportunity to explore these processes. EEG allows for precise event-based detection of neural signal, while fMRI provides insight into the broader spatial patterns of brain activation and functional connectivity (Horovitz et al., 2008; Huang et al., 2024; Laufs, 2008; Laufs, Walker, & Lund, 2007; Spoormaker et al., 2010). Previous EEG-fMRI studies on sleep have examined both global sleep characteristics (Hale et al., 2016; Moehlman et al., 2019) and the neural correlates of specific waves, including slow oscillations and spindles. These studies have generally shown that slow oscillations are associated with widespread cortical and subcortical BOLD changes (Czisch et al., 2009; Ilhan-Bayrakcı et al., 2022; Picchioni et al., 2011), whereas spindles have been linked not only to thalamic activation but also to cortical and paralimbic regions, including the hippocampus and parahippocampal gyrus (Bergmann et al., 2012; Caporro et al., 2012; Fogel et al., 2017; Schabus et al., 2007). Although these findings provide important insight into the BOLD correlates of sleep rhythms, most previous studies have focused on individual oscillatory events rather than explicitly modelling their temporal interaction (Bergmann et al., 2012; Caporro et al., 2012; Fogel et al., 2017; Picchioni et al., 2011). Only a few recent studies have begun to examine coupling between rhythms directly, for example Huang et al. (2024).

Understanding sleep rhythm-related brain activation is also essential for uncovering its potential cognitive functions. Since performing cognitive tasks during sleep is limited, most neuroimaging studies on human sleep rely on the targeted memory reactivation (TMR) paradigm, where participants are re-exposed to sensory cues during sleep to trigger associated memory reactivation (Cousins et al., 2016; Hu et al., 2020; Oudiette & Paller, 2013; Siefert et al., 2024). For instance, fMRI studies have shown significant hippocampal activation in response to odour re-exposure during sleep (Rasch et al., 2007), while EEG studies suggest that this reactivation is more likely during the UP-state of SOs (Cousins et al., 2016; Göldi et al., 2019; Xia et al., 2023). However, while TMR provides insight into externally induced memory processing during sleep, it offers limited understanding of the endogenous neural mechanisms underlying natural sleep rhythms.

To address this gap, our study investigates brain-wide activation and functional connectivity patterns associated with SO-spindle coupling, and employs a cognitive state decoding approach (Margulies et al., 2016; Yarkoni et al., 2011) - albeit indirectly - to infer potential cognitive functions. In the current study, we used simultaneous EEG-fMRI recordings during nocturnal naps (detailed sleep staging results are provided in the Methods and **Table S1**) in 107 participants. Although directly detecting hippocampal ripples using scalp EEG or fMRI is challenging, we expected that hippocampal activation in fMRI would coincide with SO-spindle coupling detected by EEG, given that SOs, spindles, and ripples frequently co-occur during NREM sleep. We also anticipated a critical role of the thalamus, particularly thalamic spindles, in coordinating hippocampal-cortical communication.

We found significant coupling between SOs and spindles during NREM sleep (N2/3), with spindle peaks occurring slightly before the SO peak. This coupling was associated with increased activation in both the thalamus and hippocampus, with functional connectivity patterns suggesting thalamic coordination of hippocampal-cortical communication, in line with prior EEG-fMRI studies of spindle-related activity (Bergmann et al., 2012; Caporro et al., 2012; Fogel et al., 2017; Schabus et al., 2007). These findings highlight the key role of the thalamus in coordinating hippocampal-cortical interactions during human sleep and provide new insights into the neural mechanisms underlying sleep-dependent brain communication. A deeper understanding of these mechanisms may contribute to future neuromodulation approaches aimed at enhancing sleep-dependent cognitive function and treating sleep-related disorders.

## Results

Simultaneous EEG-fMRI data during nocturnal sleep were collected from 138 healthy subjects (age, 22.56 ± 0.30; 81 females). After excluding subjects due to excessive head motion (mean framewise displacement > 0.5) or insufficient scanning sleep duration (<10 min), 107 participants (age, 22.40 ± 0.34; 63 females) were retained for subsequent analysis, with an average scanning time of sleep of 3.01 ± 0.14 hours (see Methods and **Table S1**). Since NREM sleep predominantly occurs in early sleep (Carskadon & Dement, 2005), we conducted our scans during the first half of the night to ensure sufficient coverage of NREM sleep, this is also the period when sleep rhythms are most consistently observed and studied in relation to memory consolidation (Maingret et al., 2016; Molle et al., 2011; Schreiner et al., 2021; Staresina et al., 2015; Staresina et al., 2023).

### Sleep stages and sleep rhythms

The primary objective of the EEG analysis was to examine the intrinsic characteristics of sleep rhythms and to identify the timing of these SO and spindle events for subsequent fMRI analyses.

We first removed MRI gradient noise from the EEG signal (detailed in Methods, see **Fig. S1**) and then applied an automated sleep staging algorithm (Vallat & Walker, 2021). The staging results were manually reviewed by two sleep experts to ensure the accurate classification of each sleep stage (**Fig. 1b**). Upon validation, we confirmed that N2/3 stages predominated in the dataset, with a significantly higher proportion of N2/3 sleep (75.77% ± 1.50%) compared to N1 (14.57% ± 1.06%, *t*_(106)_ = 25.04, *p* < 1e-4, paired-samples *t* test) and REM sleep (9.66% ± 0.90%, *t*_(106)_ = 29.60, *p* < 1e-4, **Fig. 1d, Table S1**).

**Fig. 1.**
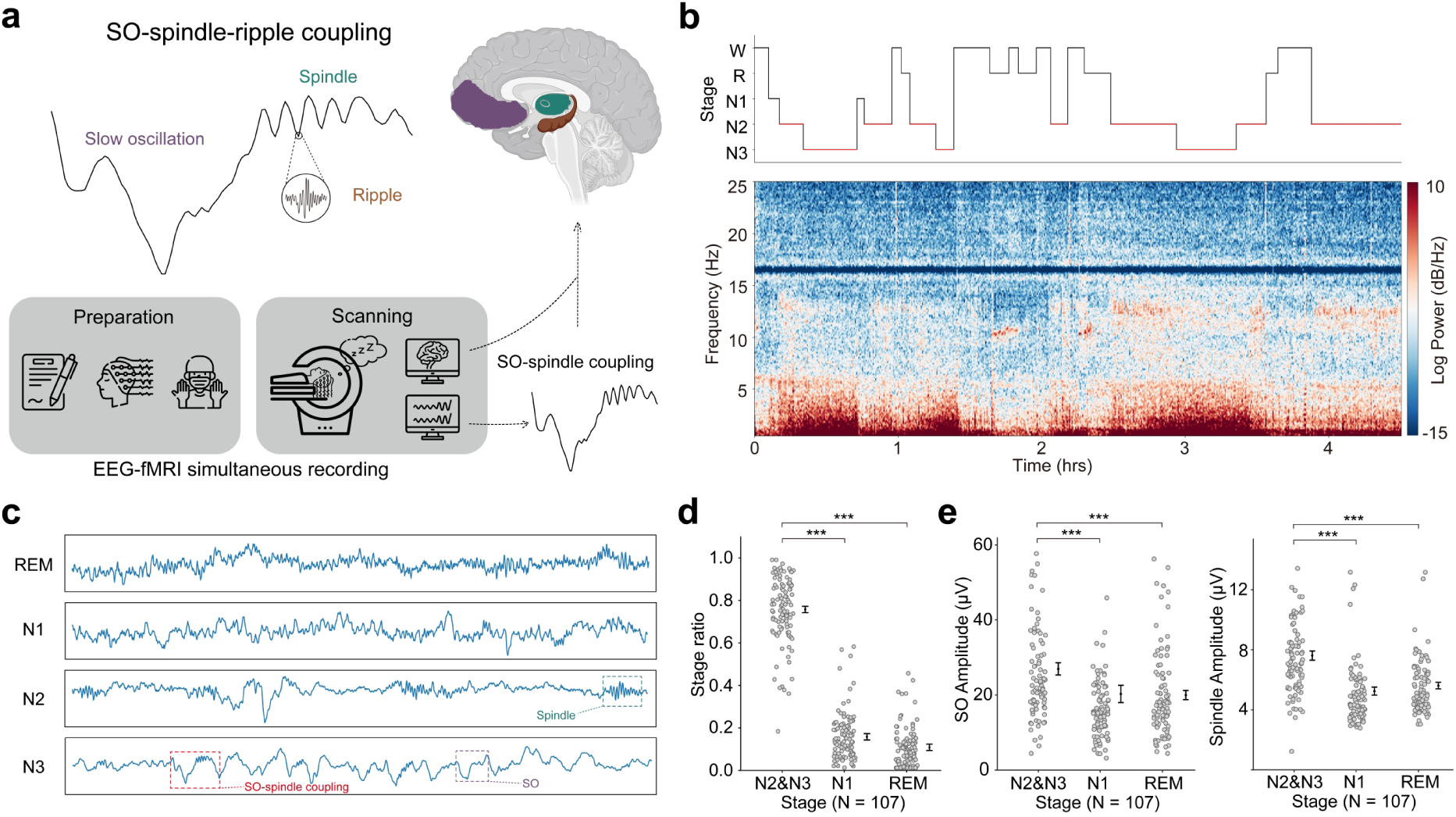
Sleep stages and sleep rhythms in 107 subjects. **a,** Sleep rhythms and task schematic. Subjects slept first half of the night with simultaneous EEG-fMRI recordings. Since detecting hippocampal ripples directly from scalp EEG is challenging, our focus was on capturing SOs, spindles, and their couplings. Regions of interest (ROIs) are colour-coded: green for the thalamus (spindle), purple for the mPFC (SOs), and orange for the hippocampus (ripples). **b,** Sleep staging and EEG spectrogram. N2/3 sleep stages (red line) were initially identified using an offline automatic sleep staging algorithm (Vallat & Walker, 2021) and then manually validated. **c,** Schematic of EEG data across different sleep stages, using preprocessed data from the C3 electrode. **d,** Proportion of each sleep stage in the dataset. **e,** Amplitudes (μV) of detected SOs (left) and spindles (right) across sleep stages. SO and spindle detection thresholds were defined from N2/3 sleep within each participant and then applied unchanged to N1 and REM for descriptive comparison. Detections in N1 and REM should therefore be interpreted as detector outputs under this fixed N2/3-derived criterion. The SO amplitudes were measured from the 0.16-1.25 Hz filtered EEG data, and spindle amplitudes were measured from the 12-16 Hz filtered EEG data. Each dot represents an individual participant. Error bars indicate SEM. *** *p* < 0.001.

Each sleep stage is characterised by distinct spectral properties and rhythmic waveforms, serving as physiological markers (**Fig. 1c**). Because SO and spindle detection relies on amplitude-based percentile thresholds, we avoided estimating separate thresholds within each sleep stage. Instead, for each participant, the SO and spindle thresholds were defined from N2/3 sleep only, where these rhythms are most abundant and physiologically expected, and the same fixed thresholds were then applied to N1 and REM for descriptive comparison.

Under this fixed N2/3-derived thresholding, detected SOs and spindles were larger and more frequent in N2/3 than in N1 or REM. SO and spindle amplitudes were significantly higher during N2/3 sleep (SO: 25.59 ± 1.49 μV; spindle: 7.39 ± 0.27 μV) than during N1 (SO: 20.15 ± 2.32 μV; spindle: 5.23 ± 0.27 μV) and REM sleep (SO: 19.84 ± 1.22 μV; spindle: 5.60 ± 0.22 μV; all *p* < 1e-4; **Fig. 1e, Fig. S2**). The corresponding event densities showed the same pattern, with 9.64 ± 0.25 SOs/min and 4.19 ± 0.10 spindles/min in N2/3, compared with 2.95 ± 0.16 SOs/min and 2.71 ± 0.14 spindles/min in N1, and 2.07 ± 0.17 SOs/min and 1.81 ± 0.14 spindles/min in REM (all *p* < 1e-4). We therefore report detections in N1 and REM only as descriptive outputs of the detector under a fixed N2/3-derived criterion, rather than as physiological equivalents of canonical N2/3 SOs or spindles.

This aligns with the established understanding that N2/3 stages feature more pronounced slow-wave activity and robust spindles (Gaillard & Blois, 1981). Furthermore, we found that the amplitude of the SO DOWN-state (13.62 ± 0.67 μV) was significantly higher than the UP-state (12.98 ± 0.66 μV, *t*_(106)_ = 4.82, *p* < 1e-4), consistent with previous work (Cash et al., 2009; Dang-Vu et al., 2008). Based on this, we used the DOWN-state to lock the timing of SOs for subsequent EEG-informed fMRI analysis. Descriptive information on sleep rhythms during these stages for all 107 subjects can be found in **Table S2-S4**.

### SO-spindle coupling

SO-spindle coupling is considered important for sleep-dependent memory consolidation. In the current study, using the same N2/3-derived detection thresholds described above, we found that SO-spindle coupling occurred most frequently during N2/3 sleep (2.46 ± 0.06 events/min). Coupling density was significantly lower in N1 (0.75 ± 0.05 events/min, *t*_(106)_ = 23.54, *p* < 1e-4) and REM sleep (0.43 ± 0.04 events/min, *t*_(106)_ = 31.24, *p* < 1e-4; **Fig. 2b, Table S2-S4**), consistent with the expected predominance of SO-spindle coupling in NREM sleep (Ngo et al., 2013; Staresina et al., 2015). As with the individual SO and spindle detections, coupling events detected in N1 and REM were retained only for descriptive stage-wise reporting (see **Table S2, S4**). They were not used to support physiological claims about SO-spindle coupling in these stages, and they were not entered into the EEG-informed fMRI analyses. All subsequent fMRI GLM and PPI analyses were restricted to N2/3 sleep.

**Fig. 2.**
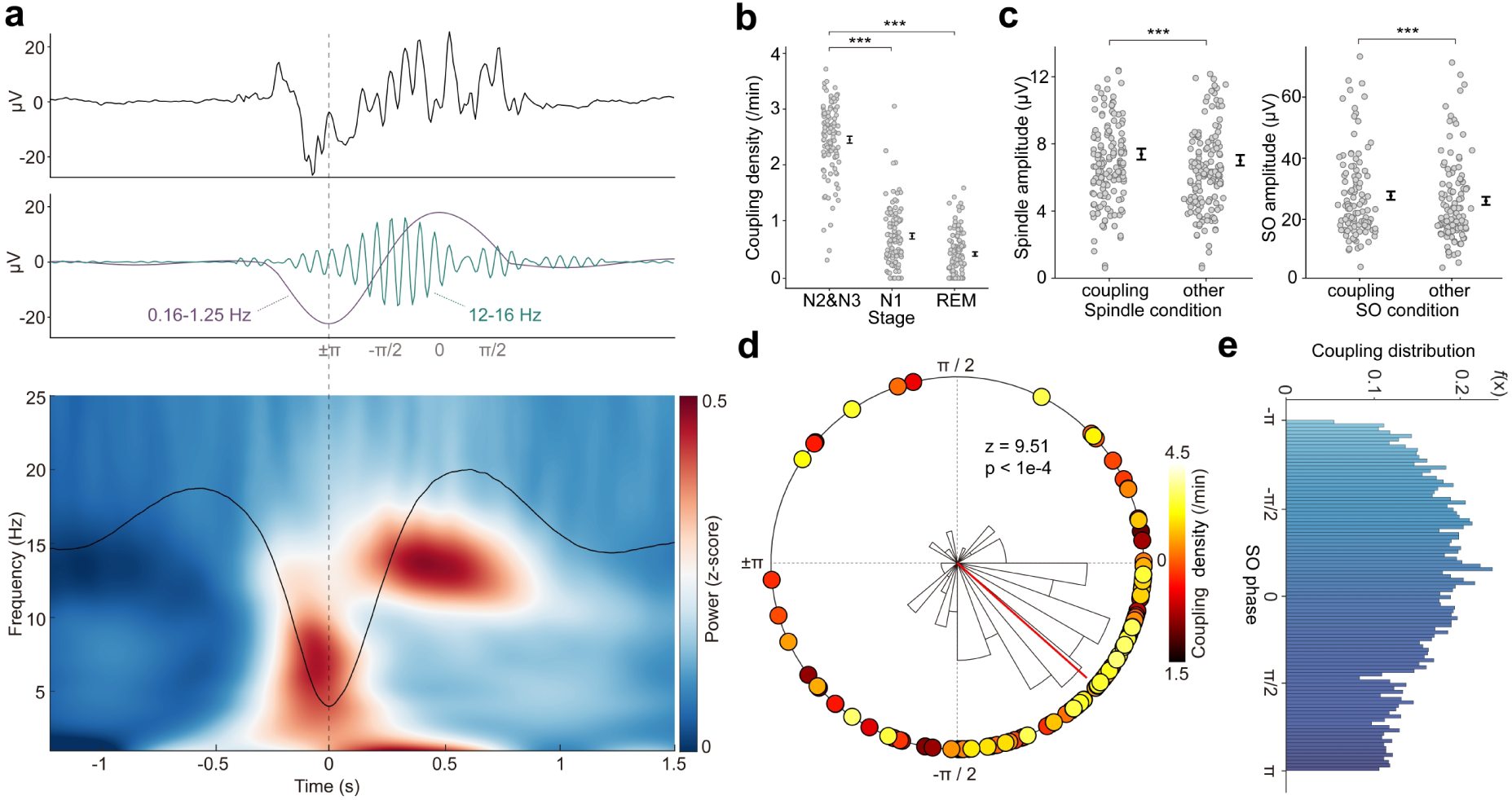
Sleep rhythms and SO-spindle coupling. **a,** The SO-spindle coupling in the temporal frequency domain. The upper two rows illustrates the spindle (12-16 Hz) phase-locked in the transition to UP-state of SO (0.16-1.25 Hz). The bottom row shows the averaged temporal frequency pattern across all instances of SO-spindle coupling and over all subjects. **b,** SO-spindle coupling density across sleep stages, using SO and spindle detections obtained with fixed N2/3-derived thresholds. Coupling events in N1 and REM are shown only for descriptive comparison. The EEG-informed fMRI analyses used N2/3 coupling events only. **c,** Differences between coupled and uncoupled sleep rhythms. The left panel shows the difference in amplitude between spindles coupled with SOs (Coupling) and spindles not coupled with SOs (Other). The right panel displays the difference in amplitude between SOs coupled with spindles (Coupling) and SOs not coupled with spindles (Other). **d,** Phase modulation of SO-spindle coupling. Spindle peaks cluster slightly before the UP-state peak of SO (i.e., 0°), where –π/2 reflects the transition from DOWN to UP-state. The histogram represents the distribution of coupling directions across all subjects, with the red line showing the mean. Coupling phases for each subject are plotted on the circle, with coupling strength colour-coded. **e,** Distribution of spindle peaks on the SO phase during all SO-spindle coupling events across participants. The distribution is represented by a probability density function, and the density is evaluated at 100 equally spaced points covering the data range. Each dot represents data from an individual subject. Error bars indicate the SEM. *** *p* < 0.001.

After extracting all N2/3 EEG epochs in which SO-spindle coupling occurred, we analysed their spectral and phase characteristics. The spindles were most likely to occur slightly before the UP-state peak of SOs (**Fig. 2a, e**), aligning with results from both animal studies (Maingret et al., 2016) and human research (Staresina et al., 2015). In our data, this pattern was consistent across subjects (**Fig. 2d**, Rayleigh test: *z* = 9.51, *p* < 1e-4), with the peak of the spindle aligned at an SO phase of −41.61 ± 0.86° (the SO UP-state peak is 0°).

Interestingly, spindle events coupled with SOs had higher amplitudes (7.69 ± 0.26 μV) compared to those not coupled (7.33 ± 0.25 μV, *t*_(106)_ = 4.65, *p* < 1e-4, **Fig. 2c**, left). Similarly, SO events coupled with spindles exhibited higher amplitudes (26.28 ± 1.34 μV) than those not coupled (24.56 ± 1.33 μV, *t*_(106)_ = 6.86, *p* < 1e-4, **Fig. 2c**, right). These findings suggest coordinated neural processing during SO-spindle coupling. Next, we explore the brain-wide activation associated with these sleep rhythms.

### Sleep rhythms associated brain activation

To investigate brain-wide activity during sleep rhythms, we modelled the fMRI blood oxygen level-dependent (BOLD) signal based on EEG-identified timing of SOs and spindle events during N2/3 sleep stages. The design matrix included the main effects of SOs, spindles, and their coupling (**Fig. 3a**, detailed in Methods). This method is validated in our previous work to effectively capture replay-aligned brain activation during rest (Huang et al., 2024). We found that SOs were associated with positive activation in the thalamus (ROI analysis, *t*_(106)_ = 2.38, *p* = 0.0096, one-sample *t* test) and significant deactivation in the neocortex (**Fig. 3b**, see also **Fig. S5**), particularly in the default mode network (DMN), which includes the mPFC and posterior cingulate cortex (PCC), consistent with previous research (Dang-Vu et al., 2008; Picchioni et al., 2011; Tagliazucchi et al., 2013). This deactivation of the neocortex, particularly the DMN, at the trough of SOs is intriguing, given that the DMN is thought to encode the internal model of the world or cognitive map (Baldassano, Hasson, & Norman, 2018; Constantinescu, O’Reilly, & Behrens, 2016; Park, Miller, & Boorman, 2021).

**Fig. 3.**
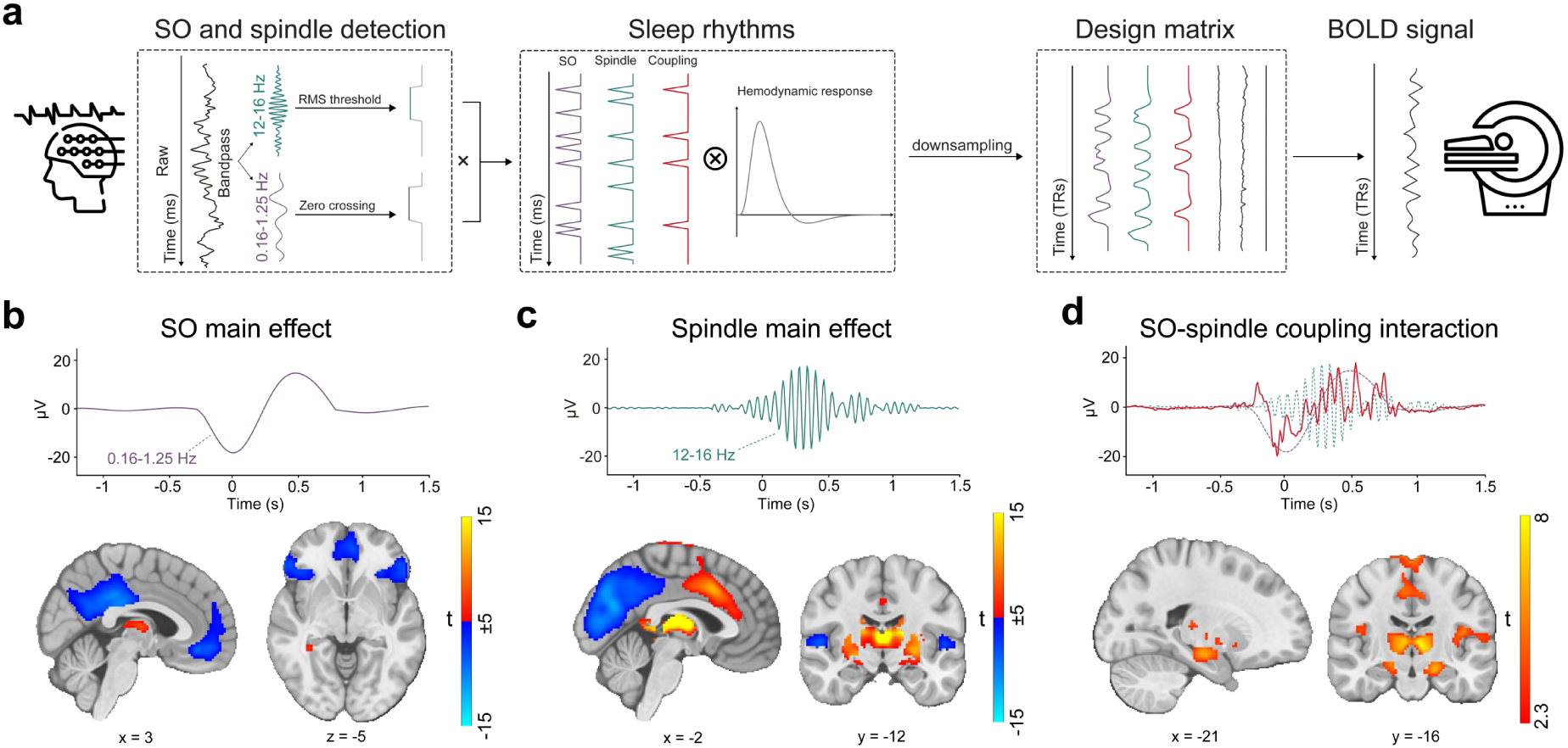
Brain-wide activation associated with sleep rhythms. **a,** Simultaneous EEG-fMRI analysis framework for detecting brain-wide activation during sleep rhythms. Detected SOs, spindles, and their coupling were convolved with the hemodynamic response function (HRF) and downsampled to match fMRI temporal resolution. These events formed the design matrix for the general linear model (GLM) analysis of fMRI activity during sleep, linking the EEG-derived timing of sleep rhythms to the corresponding brain responses in fMRI. **b,** Brain-wide activation associated with SOs. The upper row illustrates SOs, and the lower row shows the fMRI activation pattern during SO events, whole-brain family-wise error (FWE) corrected at the cluster level (*p* < 0.05) with a cluster-forming voxel threshold of *p* < 0.001. **c,** Brain-wide activation associated with spindles. Same as panel b, but for spindle events. **d,** Brain-wide activation associated with SO-spindle coupling (compared to non-coupling events).

Spindles were correlated with positive activation in the thalamus (ROI analysis, *t*_(106)_ = 15.39, *p* < 1e-4), the anterior cingulate cortex (ACC), and the putamen, alongside deactivation in the DMN **(Fig. 3c**). Notably, SO-spindle coupling was linked to significant activation in both the thalamus (ROI analysis, *t*_(106)_ = 3.38, *p* = 0.0005) and the hippocampus (ROI analysis, *t*_(106)_ = 2.50, *p* = 0.0070, **Fig. 3d**). However, no decrease in DMN activity was found during SO-spindle coupling, and DMN activity during SO was significantly lower than during coupling (ROI analysis, *t*_(106)_ = −4.17, *p* < 1e-4). For more detailed activation patterns, see **Table S5-S7**.

To ensure the results were not driven by individual differences or parameter selection, we conducted a series of control analyses. First, we excluded participants with less than 10 minutes of N2/3 sleep. Only three participants met this criterion, and their exclusion did not change the main results. For example, hippocampal activation during SO-spindle coupling remained significant (*t*_(103)_ = 2.50, *p* = 0.0071), comparable to the full sample (*t*_(106)_ = 2.50, *p* = 0.0070). Second, because the absolute number of detected SO-spindle coupling events depends on the SO detection threshold, we examined whether the main EEG-fMRI results were sensitive to this parameter. To this end, we varied the SO percentile threshold and reconstructed the EEG-informed GLM at each level. Hippocampal activation during SO-spindle coupling remained significant across a range of thresholds (71st - 80th percentile; **Fig. S6**). Third, to test whether the results depended on the use of a single lateralised frontal electrode, we repeated the EEG-informed fMRI GLM using events detected from Fz. Hippocampal activation during SO-spindle coupling again remained significant (*t*_(106)_ = 2.47, *p* = 0.0076), closely matching the original F3-based result (*t*_(106)_ = 2.50, *p* = 0.0070).

To understand the functional implications of these sleep rhythm-related brain activation, we performed open-ended cognitive state decoding using the NeuroSynth database (Yarkoni et al., 2011), examining associations between topic terms and regions of interest derived from the activation patterns of the sleep rhythms. Topic terms were ranked by their *z*-scores across the rhythms to assess overall functional relevance (see Methods for details). For positive activation patterns (**Fig. S7a**), SOs were associated with declarative (*z* = 6.31, *p* < 1e-4) and episodic memory (*z* = 5.08, *p* < 1e-4), while spindles aligned most with working memory (*z* = 4.08, *p* < 1e-4). SO-spindle coupling showed the strongest similarity to episodic (*z* = 3.66, *p* = 0.0003) and declarative memory (*z* = 2.21, *p* = 0.0271). For negative (deactivation) patterns (**Fig. S7b**), SOs were linked to declarative memory (*z* = 7.61, *p* < 1e-4), while spindles resembled task-related activations (*z* = 7.05, *p* < 1e-4). SO-spindle coupling had the highest similarity to task-related (*z* = 9.40, *p* < 1e-4) and working memory (*z* = 9.15, *p* < 1e-4). These findings suggest that internal memory-related regions are more likely to reactivate during SO-spindle coupling, whereas task-related activations, like working memory, are less prominent.

### Functional connectivity during SO-spindle coupling

Previous studies have indicated that the hippocampal-cortical dialogue during sleep memory consolidation relies on the thalamus (Coulon, Budde, & Pape, 2012; Ferraris et al., 2021; Latchoumane et al., 2017). To explore this in the human brain, we examined how functional connectivity (FC) changes when spindles couple (versus not) with SOs during sleep, using psychophysiological interaction (PPI) analysis (Friston et al., 1997). Building on the previous GLM (**Fig. 3a**), this model incorporated two additional regressors: the BOLD signal of the ROI seed and its interaction with SO-spindle events (**Fig. 4a**, detailed in Methods).

**Fig. 4.**
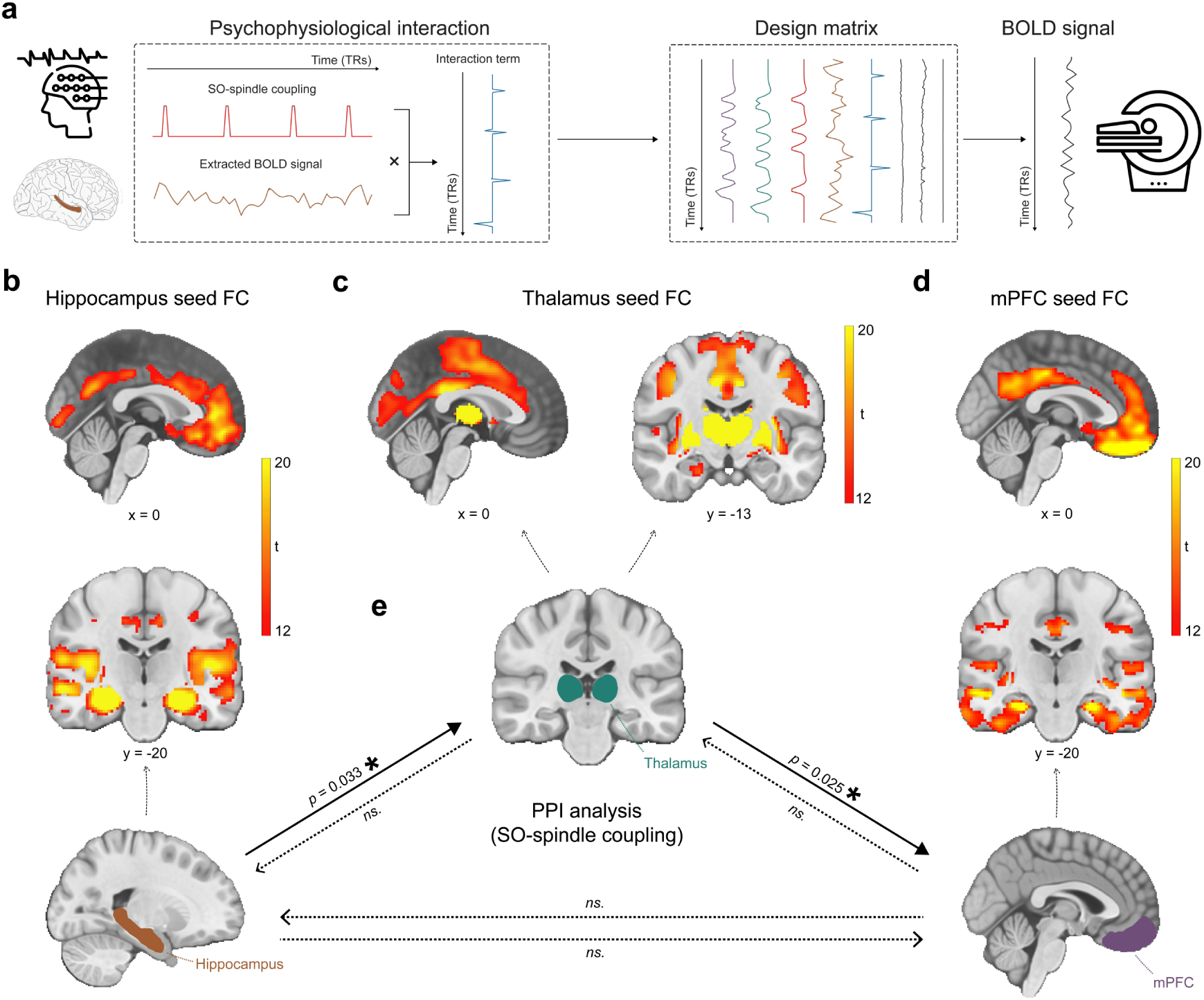
Functional connectivity changes during SO-spindle coupling. **a,** The PPI analysis framework for detecting brain-wide connectivity changes during SO-spindle coupling. This starts by setting a specific ROI (e.g., the hippocampus) as the seed to extract the BOLD signal (physiological condition) and using identified SO-spindle coupling events as the psychological condition to compute the interaction term. The design matrix includes the main effects of the physiological and psychological conditions, along with their interaction. This analysis examines whether whole-brain communication with the hippocampus changes as a function of SO-spindle coupling. **b**, Hippocampus-based functional connectivity with the whole brain (main effect of hippocampus BOLD signal in PPI analysis). The hippocampus ROI is bilateral, anatomically defined (bottom, orange colour). Brain-wide connectivity is shown with whole-brain FWE correction at the cluster level (*p* < 1e-7) with a cluster-forming voxel threshold of *p_unc._* < 0.001 for visualization purpose. **c,** Same with panel b, but based on thalamus (bilateral anatomically defined ROI). **d,** Same with panel b, but based on the mPFC (bilateral functionally defined ROI, detailed in Methods). **e,** Functional connectivity changes during SO-spindle coupling for hippocampus-based (left bottom, orange colour), thalamus-based (middle, green colour), and mPFC-based (right bottom, purple colour) connectivity. The results of ROI analysis for each direction are shown on the arrows. * *p* < 0.05, ns., not significant. Abbreviations: FC - functional connectivity, PPI - psychophysiological interaction.

We first examined whole-brain FC (physiological term) in three ROIs: the hippocampus (bilateral, anatomically defined), thalamus (bilateral, anatomically defined), and mPFC (bilateral, functionally defined). The hippocampus-based FC was mainly connected with the DMN, particularly the mPFC (**Fig. 4b**). The thalamus-based FC was primarily linked to the basal ganglia and ACC (**Fig. 4c**), and the mPFC-based FC showed strong connections with the hippocampus and PCC (**Fig. 4d**).

Crucially, the interaction between FC and SO-spindle coupling revealed that only the functional connectivity of hippocampus -> thalamus (ROI analysis, *t*_(106)_ = 1.86, *p* = 0.0328) and thalamus -> mPFC (ROI analysis, *t*_(106)_ = 1.98, *p* = 0.0251) significantly increased during SO-spindle coupling, with no significant changes in all other pathways (**Fig. 4e**). We also conducted PPI analyses for the other two events (SOs and spindles), and neither yielded significant connectivity changes in the three ROIs, as all failed to survive whole-brain FWE correction at the cluster level (*p* < 0.05). Together, these findings suggest that the thalamus, likely via spindles, coordinates hippocampal-cortical communication selectively during SO-spindle coupling, but not isolated SOs or spindle events alone.

## Discussion

Using simultaneous EEG and fMRI during nocturnal naps in 107 subjects, we identified robust coupling between spindles and the UP-state of SOs in deep NREM sleep (N2/3 stages). This coupling was linked to distinct neural activation patterns, including significant thalamic activation during spindle occurrences, reduced activity in the DMN during SOs trough, and selective hippocampal activation when spindles and SOs coupled. Furthermore, functional connectivity analyses revealed increased connectivity from the hippocampus to the thalamus and from the thalamus to the mPFC, only during SO-spindle coupling, but not isolated SOs or spindle events alone. These results shed light on how the thalamus may coordinate hippocampal-cortical interactions during NREM sleep.

Our data provide insights into the neurobiological underpinnings of these sleep rhythms. SOs, originating mainly in neocortical areas such as the mPFC, alternate between DOWN- and UP-states. The thalamus generates sleep spindles, which in turn couple with SOs. Our finding that spindle peaks consistently occurred slightly before the UP-state peak of SOs (in 83 out of 107 participants), concurs with prior studies, including Schreiner et al. (2021). Yet it differs from some results suggesting spindles might peak right at the SO UP-state (Hahn et al., 2020; Helfrich et al., 2018). Such discrepancies could arise from differences in detection algorithms, participant age (Helfrich et al., 2018), or subtle variations in cortical-thalamic timing. Nonetheless, these results underscore the importance of coordinated SO-spindle interplay in supporting sleep-dependent processes.

When modelling the timing of these sleep rhythms in the fMRI, we observed hippocampal activation selectively during SO-spindle events. This suggests the possibility of triple coupling (SOs–spindles–ripples), even though our scalp EEG was not sufficiently sensitive to detect hippocampal ripples—key markers of memory replay (Buzsáki, 2015). Recent iEEG evidence indicates that ripples often co-occur with both spindles (Ngo, Fell, & Staresina, 2020) and SOs (Staresina et al., 2015; Staresina et al., 2023). Therefore, the hippocampal involvement during SO-spindle events in our study may reflect memory replay from the hippocampus, propagated via thalamic spindles to distributed cortical regions.

The thalamus, known to generate spindles (Halassa et al., 2011), plays a key role in producing and coordinating sleep rhythms (Coulon, Budde, & Pape, 2012; Crunelli et al., 2018), while the hippocampus is found essential for memory consolidation (Buzsáki, 2015; Diba & Buzsáki, 2007; Singh, Norman, & Schapiro, 2022). The increased hippocampal and thalamic activity, along with strengthened connectivity between these regions and the mPFC during SO-spindle events, underscores a hippocampal-thalamic-neocortical information flow. This aligns with recent findings suggesting the thalamus orchestrates neocortical oscillations during sleep (Schreiner et al., 2022). The thalamus and hippocampus thus appear central to memory consolidation during sleep, guiding information transfer to the neocortex, e.g., mPFC.

An intriguing aspect of our findings is the reduced DMN activity during SOs when modelled at the SO trough (DOWN-state). This reduced DMN activity may reflect large-scale neural inhibition characteristic of the SO trough. The DMN is typically active during internally oriented cognition (e.g., self-referential processing or mind-wandering) and is suppressed during external stimuli processing (Yeshurun, Nguyen, & Hasson, 2021). It is unlikely, however, that this suppression of DMN during SO events is related to a shift from internal cognition to external responses given it is during deep sleep time. Instead, it could be driven by the inherent rhythmic pattern of SOs, which makes it difficult to separate UP- from DOWN-states (the two temporal regressors were highly correlated, and similar brain activation during SOs events was obtained if modelled at the SO peak instead, **Fig. S5**). Since the amplitude at the SO trough is consistently larger than that at the SO peak, the neural activation we detected may primarily capture the large-scale inhibition from DOWN-state. Interestingly, no such DMN reduction was found during SO-spindle coupling, implying that coupling may involve distinct neural dynamics that partially re-engage DMN-related processes, possibly reflecting memory-related reactivation. Future research using high-temporal-resolution techniques like iEEG could clarify these possibilities.

To explore functional relevance, we employed an open-ended cognitive state decoding approach using meta-analytic data (NeuroSynth: Yarkoni et al. (2011)). Although this method usefully generates hypotheses about potential cognitive processes, particularly in the absence of a pre- and post-sleep memory task, it is inherently indirect. Many cognitive terms showed significant associations (16 of 50), such as “episodic memory,” “declarative memory,” and “working memory.” We focused on episodic/declarative memory given the known link with hippocampal reactivation (Diekelmann & Born, 2010; Staresina et al., 2015; Staresina et al., 2023). Nonetheless, these inferences regarding memory reactivation should be interpreted cautiously without direct behavioral measures. Future research incorporating explicit tasks before and after sleep would more rigorously validate these potential functional claims.

Despite providing new insights, our study has several limitations. First, our scalp EEG did not directly capture hippocampal ripples, preventing us from conclusively demonstrating triple coupling. Second, the combination of EEG-fMRI and the lack of a memory task limit our ability to parse fine-grained BOLD responses at the DOWN- vs. UP-states of SOs and link observed activations to behavioral outcomes. Third, sleep oscillation detection was based on a single frontal electrode. This choice improved signal stability and event timing in the prolonged simultaneous EEG-fMRI setting, but it did not exploit the full multichannel EEG information and cannot characterise the full spatial distribution of SOs and spindles. In particular, F3-based detection may be more sensitive to frontal SOs and frontal sigma activity than to the centro-parietal fast spindle component. We therefore interpret the EEG-informed fMRI results as reflecting BOLD activity associated with frontal-channel detected SOs, spindles, and their coupling during N2/3 sleep. Future studies using multichannel or source-informed detection strategies, with separate treatment of slow and fast spindles, will be better suited to capture the spatial dynamics of these sleep oscillations. Fourth, the use of large anatomical ROIs may mask subregional contributions of specific thalamic nuclei or hippocampal subfields. Finally, without a memory task, we cannot establish a direct behavioral link between sleep-rhythm-locked activation and memory consolidation. Future studies combining ultra-high-field fMRI or iEEG with cognitive tasks, as well as multichannel or source-informed detection strategies that separately characterize slow and fast spindles, will be better suited to refine our understanding of subregional network dynamics and the functional significance of sleep oscillations.

In summary, we found that SO-spindle coupling during deep NREM sleep is associated with distinct activation in both the thalamus and hippocampus, accompanied by increased connectivity from the hippocampus to the thalamus and from the thalamus to the mPFC. These results emphasize the central role of the thalamus in orchestrating hippocampal-cortical interactions and highlight potential neural substrates underlying the complex interplay of SOs, spindles, and, possibly, ripples. Our open-ended decoding analyses suggest that these coordinated sleep events may be relevant for memory-related processes; however, future experiments incorporating explicit behavioral measures are necessary to confirm their functional significance. By revealing how these rhythms modulate widespread brain networks, our findings open avenues for developing targeted neuromodulatory strategies to enhance sleep-dependent cognitive functions and potentially address sleep-related disorders.

## Methods

### Participants & Protocol

A total of 138 healthy adults (age: 22.56 ± 0.30; 81 females) were recruited for the study. All participants had either normal vision or vision corrected to normal standards. None had any past psychiatric or neurological disorders. Prior to participation, they were screened for eligibility to undergo magnetic resonance imaging (MRI). The experiment received approval from Peking University ethics committee (reference number: #2015-26, #2018-11-05). Every participant provided written informed consent.

At the start of the experiment, subjects arrived early at the laboratory for registration. They signed informed consent forms, washed their hair, changed their clothes, and put on the EEG cap in preparation for the experiment. After these preliminary steps, participants entered the MRI scanning room by around 20:30, settled down for sleep, and data collection began. Given concern for potential data drift and decline in the signal-to-noise ratio due to extended and continuous fMRI scanning, the data acquisition was segmented into several sessions for data collection. A “sleep” session was concluded either when a participant was fully awake (unable to fall asleep again or needed to use the restroom) or after 8192 seconds (the maximum scanning time of the scan, equating to 4096 volumes with 2 seconds TR).

After removing subjects with excessive head movements (mean FD > 0.5) or those whose sleep durations were under 10 minutes, the final participant count for further analyses was 107 (age: 22.40 ± 0.34; 63 females) with a mean scanning duration of sleep of 3.01 ± 0.14 hours.

### EEG data acquisition

EEG was recorded simultaneously with fMRI data using an MR-compatible EEG amplifier system (BrainAmps MR-Plus, Brain Products, Germany), along with a specialized electrode cap. The recording was done using 64 channels in the international 10/20 system, with the reference channel positioned at FCz. In order to adhere to polysomnography (PSG) recording standards, six electrodes were removed from the EEG cap: one for electrocardiogram (ECG) recording, two for electrooculogram (EOG) recording, and three for electromyogram (EMG) recording. EEG data was recorded at a sample rate of 5000 Hz, the resistance of the reference and ground channels was kept below 10 kΩ, and the resistance of the other channels was kept below 20 kΩ. To synchronize the EEG and fMRI recordings, the BrainVision recording software (BrainProducts, Germany) was utilized to capture triggers from the MRI scanner. The EEG sampling was synchronized to the MR scanner’s 10 MHz gradient system clock, ensuring a stable gradient artifact shape over time and enabling accurate artifact removal. This was achieved via the standard clock synchronization interface of the EEG amplifier, minimizing timing jitter and drift.

### MRI data acquisition

All MRI data were acquired using a 20-channel head coil on a research-dedicated 3-Tesla Siemens Magnetom Prisma MRI scanner. Earplugs and cushions were provided for noise protection and head motion restriction. We chose the 20-channel head coil because it was compatible with our EEG system and ensured sufficient signal-to-noise ratio (SNR) for both EEG and fMRI acquisition. The EEG electrode cables were guided through the lateral and posterior openings of the head coil, secured with foam padding to reduce motion and minimize MR-related artifacts. Moreover, given the extended nature of nocturnal sleep recordings, the 20-channel coil helped maintain participant comfort while still achieving high-quality simultaneous EEG-fMRI data.

For registration purposes, high-resolution T1-weighted anatomical images were acquired using a three-dimensional magnetization prepared rapid acquisition gradient-echo sequence (sequence specification: number of slices = 192, repetition time (TR) = 2,530 ms, echo time (TE) = 2.98 ms, inversion time = 1,100 ms, voxel size = 0.5 × 0.5 × 1 mm^3^, flip angle (FA) = 7°). The participants were asked to lie quietly in the scanner during data acquisition.

Then, the “sleep” session began after the participants were instructed to try and fall asleep. For the functional scans, whole-brain images were acquired using a T2*-weighted gradient echo-planar imaging (EPI) sequence sensitive to the BOLD contrast. The sequence parameters were as follows: 33 slices in interleaved ascending order, TR = 2000 ms, TE = 30 ms, voxel size = 3.5 × 3.5 × 4.2 mm^3^, FA = 90°, matrix = 64 × 64, gap = 0.7 mm. A relatively large voxel size was chosen to preserve signal-to-noise ratio while maintaining whole-brain coverage within a feasible repetition time. This compromise was important for the prolonged overnight EEG-fMRI sleep protocol, where head motion, physiological noise, participant comfort, and sustained acquisition stability are substantial practical constraints (Bodurka et al., 2007; Laufs et al., 2008). A smaller voxel size would have improved spatial specificity, but would have required either a longer repetition time, reduced brain coverage, or lower signal-to-noise ratio.

### EEG data preprocessing and sleep stage scoring

EEG data collected inside MRI scanner was contaminated by imaging, ballistocardiographic (BCG) and ocular artifacts. To deal with these artifacts, the preprocessing processes was divided into two parts: automated processing and manual processing.

The automated processing was carried out using custom Python scripts and the MNE toolkit (Gramfort et al., 2013). This encompassed the following procedures: **I**. The raw EEG data were downsampled from 5000 Hz to 2048 Hz. This downsampled rate accelerates subsequent analyses while retaining the necessary precision to distinguish MR noise. **II.** Imaging artifacts were removed using the Average Artefact Subtraction method (Allen, Josephs, & Turner, 2000). **III.** We then applied 0.1 - 30 Hz band-pass finite impulse response (FIR) filters to all channels. The EEG and EOG electrodes were re-referenced by subtracting the value from the M1 or M2 electrode (whichever was from the opposite hemisphere). The outer EMG electrodes were re-referenced to the middle one. **IV.** The signal space projection algorithm was applied to extract all epochs based on TRs and remove residual MR noise, as well as automatically extracting all heartbeat epochs using ECG electrode data for the removal of BCG noise. **V.** The data were further downsampled to 100 Hz and subjected to Independent Component Analysis (ICA).

Following this, the manual processing is carried out with labelling and removing the physiological artifacts ICs belonging to eye, muscle and residual BCG through the use of custom MATLAB scripts and the EEGLAB toolkit (Delorme & Makeig, 2004). Sleep stages were then scored by YASA toolkit (Vallat & Walker, 2021) after preprocessing, necessitating data from electrodes C3, left EOG, and left EMG. The classification of N2/3 stages were further verified by experts for validity.

### MRI data preprocessing

Results included in this manuscript come from preprocessing performed using *fMRIPrep* 21.0.4 (Esteban et al., 2020; Esteban et al., 2019), which is based on *Nipype* 1.6.1 (Gorgolewski et al., 2011). Many internal operations of *fMRIPrep* use *Nilearn* 0.8.1 (Abraham et al., 2014), mostly within the functional processing workflow. For more details of the pipeline, see https://fmriprep.readthedocs.io/en/latest/workflows.html.

*Conversion of data to the brain imaging data structure standard.* To facilitate further analysis and sharing of data, all study data were arranged according to the Brain Imaging Data Structure (BIDS) specification using dcm2bids tool, which is freely available from https://unfmontreal.github.io/Dcm2Bids/.

*Anatomical data preprocessing.* The T1-weighted (T1w) image was corrected for intensity non-uniformity (INU) with N4BiasFieldCorrection (Tustison et al., 2010), distributed with ANTs 2.3.3 (Avants et al., 2008), and used as T1w-reference throughout the workflow. The T1w-reference was then skull-stripped with a *Nipype* implementation of the antsBrainExtraction.sh workflow (from ANTs), using OASIS30ANTs as target template. Brain tissue segmentation of cerebrospinal fluid (CSF), white-matter (WM) and gray-matter (GM) was performed on the brain-extracted T1w using fast (FSL 6.0.5.1:57b01774, RRID:SCR_002823) (Zhang, Brady, & Smith, 2001). Volume-based spatial normalization to one standard space (MNI152NLin2009cAsym) was performed through nonlinear registration with antsRegistration (ANTs 2.3.3), using brain-extracted versions of both T1w reference and the T1w template. The following template was selected for spatial normalization: *ICBM 152 Nonlinear Asymmetrical template version 2009c* (Fonov, 2009) [RRID:SCR_008796; TemplateFlow ID: MNI152NLin2009cAsym].

*Functional data preprocessing.* First, a reference volume and its skull-stripped version were generated using a custom methodology of *fMRIPrep*. Head-motion parameters with respect to the BOLD reference (transformation matrices, and six corresponding rotation and translation parameters) are estimated before any spatiotemporal filtering using mcflirt (Jenkinson et al., 2002) [FSL 6.0.5.1:57b01774]. BOLD runs were slice-time corrected to 0.96s (0.5 of slice acquisition range 0s-1.92s) using 3dTshift from AFNI (Cox & Hyde, 1997). The BOLD time-series (including slice-timing correction when applied) were resampled onto their original, native space by applying the transforms to correct for head-motion. These resampled BOLD time-series will be referred to as *preprocessed BOLD in original space*, or just *preprocessed BOLD*. The BOLD reference was then co-registered to the T1w reference using mri_coreg (FreeSurfer) followed by flirt (Jenkinson & Smith, 2001) [FSL 6.0.5.1:57b01774] with the boundary-based registration (Greve & Fischl, 2009) cost-function. Co-registration was configured with six degrees of freedom. Several confounding time-series were calculated based on the *preprocessed BOLD*: framewise displacement (FD), DVARS and three region-wise global signals. FD was computed using two formulations following Power (absolute sum of relative motions) (Power et al., 2014) and Jenkinson (relative root mean square displacement between affines, Jenkinson et al. (2002)). FD and DVARS are calculated for each functional run, both using their implementations in *Nipype* (following the definitions by Power et al. (2014)). The three global signals are extracted within the CSF, the WM, and the whole-brain masks. Additionally, a set of physiological regressors were extracted to allow for component-based noise correction (*CompCor)* (Behzadi et al., 2007). Principal components are estimated after high-pass filtering the *preprocessed BOLD* time-series (using a discrete cosine filter with 128s cut-off) for the two *CompCor* variants: temporal (tCompCor) and anatomical (aCompCor). tCompCor components are then calculated from the top 2% variable voxels within the brain mask. For aCompCor, three probabilistic masks (CSF, WM and combined CSF+WM) are generated in anatomical space. The implementation differs from that of Behzadi et al. (2007) in that instead of eroding the masks by 2 pixels on BOLD space, the aCompCor masks are subtracted a mask of pixels that likely contain a volume fraction of GM. This mask is obtained by thresholding the corresponding partial volume map at 0.05, and it ensures components are not extracted from voxels containing a minimal fraction of GM. Finally, these masks are resampled into BOLD space and binarized by thresholding at 0.99 (as in the original implementation). Components are also calculated separately within the WM and CSF masks. For each CompCor decomposition, the *k* components with the largest singular values are retained, such that the retained components’ time series are sufficient to explain 50 percent of variance across the nuisance mask (CSF, WM, combined, or temporal). The remaining components are dropped from consideration. Frames that exceeded a threshold of 0.5 mm FD or 1.5 standardised DVARS were annotated as motion outliers.

The BOLD time-series were resampled into standard space, generating a *preprocessed BOLD run in MNI152NLin2009cAsym space*. First, a reference volume and its skull-stripped version were generated using a custom methodology of *fMRIPrep*. All resamplings can be performed with *a single interpolation step* by composing all the pertinent transformations (i.e., head-motion transform matrices, susceptibility distortion correction when available, and co-registrations to anatomical and output spaces). Gridded (volumetric) resamplings were performed using antsApplyTransforms (ANTs), configured with Lanczos interpolation to minimize the smoothing effects of other kernels (Lanczos, 1964). Non-gridded (surface) resamplings were performed using mri_vol2surf (FreeSurfer).

### EEG sleep rhythms analysis

In line with other studies examining the coordination between SOs and spindles, particularly their spatial distribution (Massimini et al., 2004; Molle et al., 2011), EEG data from the F3 electrode were used to quantify the occurrence of SOs and spindles. Using the established detection algorithms (detailed below), we identified SOs and sleep spindles for each subject.

It is worth noting that the primary aim of EEG rhythm detection was to identify reliable event times for EEG-informed fMRI modelling. Detection was performed on the F3 electrode because this channel provided stable signal quality during prolonged nocturnal EEG-fMRI recordings. In our MR-compatible EEG setup, FCz was used as the online reference. Central electrodes close to this reference, and electrodes near the vertex that were in prolonged contact with the head coil during supine sleep, were more susceptible to reduced signal contrast, impedance drift, and pressure-related degradation of electrode-scalp contact. We therefore used F3 as a pragmatic choice to maximize reliable event timing in N2/3 sleep. This choice was not intended to characterize the full scalp topography of SOs or spindles, and it may underrepresent the centro-parietal fast spindle component. As a sensitivity analysis, we repeated the main EEG-informed fMRI GLM using Fz, a midline frontal electrode, with the same detection and modelling procedure.

*Detection of SOs.* Data were first bandpass-filtered between 0.16 and 1.25 Hz (Butterworth filter, order 3, bidirectional filtering for zero phase). After identifying all positive-to-negative zero crossings, potential SOs were defined based on the interval between consecutive zero crossings, ranging from 0.8 s to 3 s. For each potential SO, we calculated the amplitude range as the peak minus the trough.

For each participant, the amplitude threshold was defined as the 75th percentile of candidate SO amplitude ranges observed during N2/3 sleep. This fixed N2/3-derived threshold was then applied unchanged across the recording for descriptive stage-wise summaries. Detected events were assigned to N1, N2/3 or REM according to the sleep-stage label at the event time. Only candidates exceeding this threshold were labelled as SOs, following previous work (Schreiner et al., 2021).

*Detection of sleep spindles.* Detection of sleep spindles. Data were bandpass-filtered between 12 and 16 Hz (Butterworth filter, order 3, bidirectional filtering for zero phase). The root mean square (RMS) of the filtered signal was computed with a 200 ms sliding time window. For each participant, the spindle threshold was defined as the 75th percentile of RMS values observed during N2/3 sleep. This fixed N2/3-derived threshold was then applied unchanged across the recording for descriptive stage-wise summaries. Detected events were assigned to N1, N2/3 or REM according to the sleep-stage label at the event time. RMS segments exceeding this threshold for 0.5 s to 3 s were identified as spindles (Staresina et al., 2015).

*Detection of SO-spindle couplings.* From the detected SOs and spindles, we identified the peak time of each spindle. Within each SO interval, we checked whether a spindle peak occurred; if so, that SO was labelled as an SO-spindle coupling event. For descriptive stage-wise summaries, coupling events were assigned to the sleep stage of the corresponding SO trough. For every SO-spindle coupling event, an epoch was created time-locked to the SO trough as the central reference, following Schreiner et al. (2021). We extracted data in a [−4 s to 4 s] window around this point, forming the epoch for each coupling event. For the EEG-informed fMRI analyses, only SO, spindle and SO-spindle coupling events detected during N2/3 sleep were used.

The detection procedures described above were developed primarily for N2 and N3 sleep, where SOs, spindles and their coupling are physiologically expected and most reliably observed (Hahn et al., 2020; Helfrich et al., 2019; Helfrich et al., 2018; Ngo, Fell, & Staresina, 2020; Schreiner et al., 2022; Schreiner et al., 2021; Staresina et al., 2015; Staresina et al., 2023). Because percentile-based thresholds can otherwise force the detector to label events in every sleep stage, we did not estimate separate thresholds within N1 or REM. Instead, for each participant, all SO and spindle thresholds were defined from N2/3 sleep and then applied uniformly across the recording. **Tables S1 and S3** report detailed statistical information on sleep rhythm and N2/3 events detection. The N1 and REM events detection reported in **Tables S2 and S4**, and illustrated in **Fig. S2-S4**, should therefore be interpreted as descriptive detector outputs under this fixed N2/3-derived criterion, rather than as evidence for canonical N2/3 SOs, spindles or physiological SO-spindle complexes in those stages. These detections were not used in the EEG-informed fMRI GLM or PPI analyses, which were restricted to N2/3 sleep.

*Event-related potentials (ERP) analysis.* After completing the detection of each sleep rhythm event, we performed ERP analyses for SOs, spindles, and coupling events in different sleep stages. Specifically, for SO events, we took the trough of the DOWN-state of each SO as the zero-time point, then extracted data in a [-2 s to 2 s] window from the broadband (0.1–30 Hz) EEG and used [−2 s to −0.5 s] for baseline correction; the results were then averaged across 107 subjects (see **Fig. S2a**). For spindle events, we used the peak of each spindle as the zero-time point and applied the same data extraction window and baseline correction before averaging across 107 subjects (see **Fig. S2b**). Finally, for SO-spindle coupling events, we followed the same procedure used for SO events (see **Fig. 2a**, **Figs. S3–S4**).

*Time-frequency analysis.* We performed this analysis on the SO-spindle coupling epochs using the FieldTrip toolbox (Oostenveld et al., 2011). Fourier analysis was executed on the data, utilizing a sliding time window with a forward shift of 50 ms. The window length was defined as five cycles of the present frequency (frequency range: 1-30 Hz, step size: 1 Hz). Before Fourier analysis, windowed data segments underwent multiplication by a Hanning taper. Subsequently, power values were normalized to z-scores within the time range of [-4 s to 4 s], selecting a longer duration to prevent edge artifacts from low-frequency activity in the target time window of [-1.5 s to 1.5 s].

*Preferred phase analysis.* All SO-spindle coupling epochs were filtered to the SO frequency band (0.16-1.25 Hz, 3rd-order Butterworth filter), and the instantaneous SO phase was determined through the Hilbert transform. Using the saved spindle peak times for each SO-spindle coupling, the preferred phase was deduced. The distribution of these phases across participants was then tested for uniformity using the Rayleigh test.

### fMRI GLM analysis

All fMRI GLM analyses were conducted using only data from the N2/3 stages of NREM sleep. Segments of sleep data from these stages were concatenated for each participant. After pre-processing, the fMRI data was smoothed with an 8 mm FWHM kernel. All images underwent a temporal high-pass filter (width of 100 s), and the autocorrelation of the hemodynamic response was modelled via an AR(1) model. We then conducted General Linear Model (GLM) analyses to identify whole-brain activations associated with sleep rhythms and their coupling. We incorporated nuisance regressors estimated during the preprocessing with fMRIprep: six rigid-body motion-correction parameters identified during realignment (comprising three translation and rotation parameters), the mean White Matter, and the mean Cerebral Spinal Fluid signals. The cosine drift model was employed to detect and eliminate data fluctuations, thereby enhancing the accuracy of actual experimental effects. The GLMs were designed to evaluate the influence of experimental conditions on BOLD responses. All whole-brain analyses were thresholded using whole-brain FWE corrected at the cluster level (*p* < 0.05), with a cluster-forming voxel threshold of *p*_unc._ < 0.001, unless otherwise specified.

The fMRI GLM analysis was designed to look for brain activations pertinent to SO, spindle, and SO-spindle coupling. Drawing from the EEG results concerning the timing of SO and spindle, we identified the trough of the SO as the onset for SO events and the peak of the spindle as the onset for spindle events. We chose the trough of the SO and the peak of the spindle as the onset points, primarily due to the larger amplitudes observed at these moments (thereby more robust to noise). This was also the common choice in previous EEG studies on SO and spindles (Ngo, Fell, & Staresina, 2020; Staresina et al., 2015; Staresina et al., 2023). To model SO-spindle coupling in fMRI, we followed Schreiner et al. (2021), and defining SO-spindle complex as the peak of spindle appeared within 1.5 seconds after SO trough. When such a peak was found, that particular SO was labelled as an SO-spindle event, with its SO trough labelled as the event onset. After convolving these events with the canonical Hemodynamic Response Function (HRF) of SPM and downsampling them to align with the fMRI time resolution, they were integrated as regressors in the design matrix for GLM analysis, allowing us to determine the effects of SO and spindle, as well as the SO-spindle coupling (**Fig. 3b-d**).

We have also conducted a control analysis aimed to distinguish brain activations associated with DOWN- vs. UP state of SOs. Drawing again from EEG data on SO and spindle, we randomly selected half of the SO events using the peak as the onset and the other half using the trough, to reduce collinearity between the two regressors. As with the main analysis, we identified the spindle peak as the onset for spindle events (**Fig. S5**).

### ROI analysis

The objective of our ROI analysis was to test the activation of specific brain regions during distinct sleep rhythms. Within each ROI, the beta values at the subject-level were averaged across all voxels to facilitate subsequent statistical inferences (one-sample *t* test). We defined the mDMN and mPFC ROIs functionally based on the meta-analysis results in Neurosynth (Yarkoni et al., 2011), by the keyword “medial default mode network” and “ventromedial prefrontal cortex” respectively. In addition to mDMN and mPFC, we defined other ROIs anatomically, using the high-resolution Harvard-Oxford Atlas probabilistic atlas (Desikan et al., 2006).

### Open-ended Cognitive State Functional Decoding

For functional decoding, we followed the approach in Margulies et al. (2016). We used the NeuroSynth (Yarkoni et al., 2011) meta-analytic database (www.neurosynth.org) to assess topic terms associated with the sleep rhythms activation patterns. ROI masks were created from the whole-brain activation patterns associated with the three sleep rhythms in **Fig. 3b-d**. These activation patterns were binarized using a threshold of *p* < 0.001, forming a brain mask for each sleep rhythm.

For each ROI mask, the analysis output was a z-statistic associated with a feature term, and the terms were ranked based on the weighted sum of their z-scores. The feature terms were derived from the most general set of 50 topic terms available in the NeuroSynth v3 dataset (https://github.com/NeuroanatomyAndConnectivity/gradient_analysis/blob/master/gradient_data/neurosynth/v3-topics-50-keys.txt).

From the 50 topics, 16 exceeded the threshold of *z* > 1.96 (*p* < 0.05). Two terms were removed as they did not represent coherent cognitive functions, leaving 14 relevant topic terms (**Fig. S7**). These included: episodic memory, declarative memory, working memory, task representation, language, learning, faces, visuospatial processing, category recognition, cognitive control, reading, cued attention, inhibition, and action.

### PPI analysis

PPI (psychophysiological interaction) analysis is designed to quantify context-dependent alterations in effective connectivity between brain regions. For instance, it can determine whether hippocampus-based functional connectivity with the thalamus varies based on the occurrence of an SO-spindle coupling event. We conducted nine whole-brain PPI analyses with *nilearn* to investigate interactions based in the hippocampus, thalamus or mPFC during SO-spindle coupling, isolated SO and isolated spindle, utilizing the same ROIs. Both the main effects of physiological and physiological condition, as well as their interaction were modelled in the design matrix for PPI analysis. The main effect of physiological term produced functional connectivity maps of the entire brain in relation to the specified ROI (**Fig. 4b-d**). While the interaction term asked for the *change* of functional connectivity along with the SO-spindle coupling (**Fig. 4e**).

## Acknowledgment

Author contributions: Conceptualization, Y.L., H.W., J.G.; Investigation, H.W., Q.Z., J.Z., Y.L., J.G.; Writing – Original Draft, H.W., Y.L.; Writing – Review & Editing, H.W., Y.L., J.Z., Q.Z., J.G. Funding: This study is supported by the National Science and Technology Innovation 2030 Major Program (2022ZD0205500), the National Natural Science Foundation of China (32271093), the Beijing Municipal Science and Technology Commission (Z230010, L222033), Beijing United Imaging Research Institute of Intelligent Imaging Foundation (CRIBJZD202101), the Open Research Fund of the National Center for Protein Sciences at Peking University, and the Fundamental Research Funds for the Central Universities.

## Competing interests

The authors declare that they have no competing interests.

## Data and materials availability

The data that support the findings can be provided by the corresponding author Y.L. pending scientific review and a completed material transfer agreement. Requests for the data should be submitted to: https://datadryad.org/stash/dataset/doi:10.5061/dryad.2fqz612x0. The analysis code will be publicly available on https://gitlab.com/liu_lab/eeg-fmri-sleep upon publication.

## Supplementary Materials

**Fig. S1.**
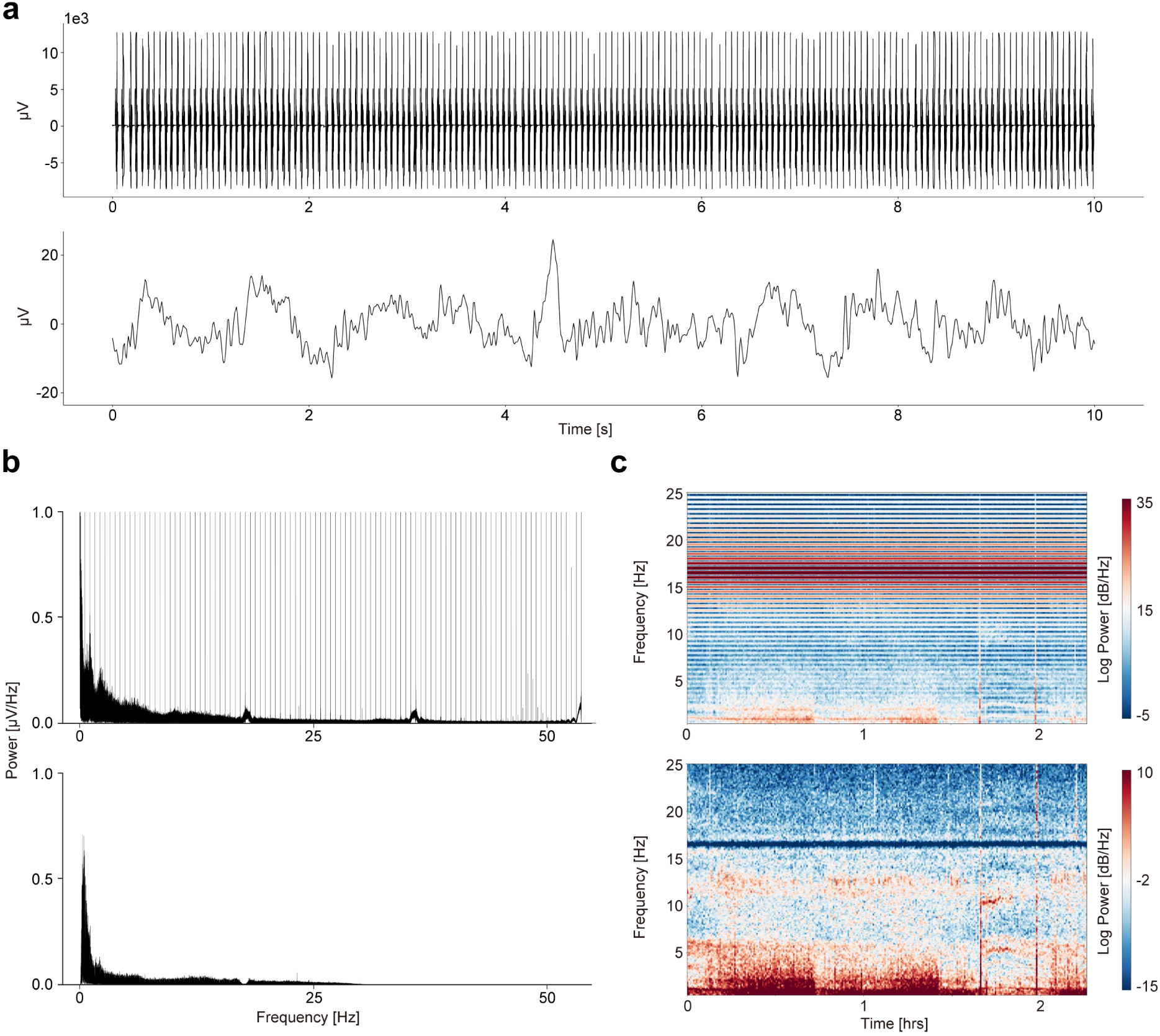
Removal of MRI gradient noise from simultaneous collected EEG data. **a**, Time series of both raw and preprocessed EEG data. The top row depicts the raw EEG data, which contains noise primarily from the MRI gradient magnetic field and electrocardiographic artifacts. The bottom row showcases the preprocessed EEG data (detailed in Methods). **b**, Power spectral density of the raw and preprocessed EEG data estimated by the fast Fourier transform. The raw EEG data is shown in the top row, while the preprocessed EEG data is in the bottom row. **c**, Time-frequency spectrogram of the raw and preprocessed EEG data, by the short-time Fourier transform. The top row represents the raw EEG data, and the bottom row displays the preprocessed EEG data.

**Fig. S2.**
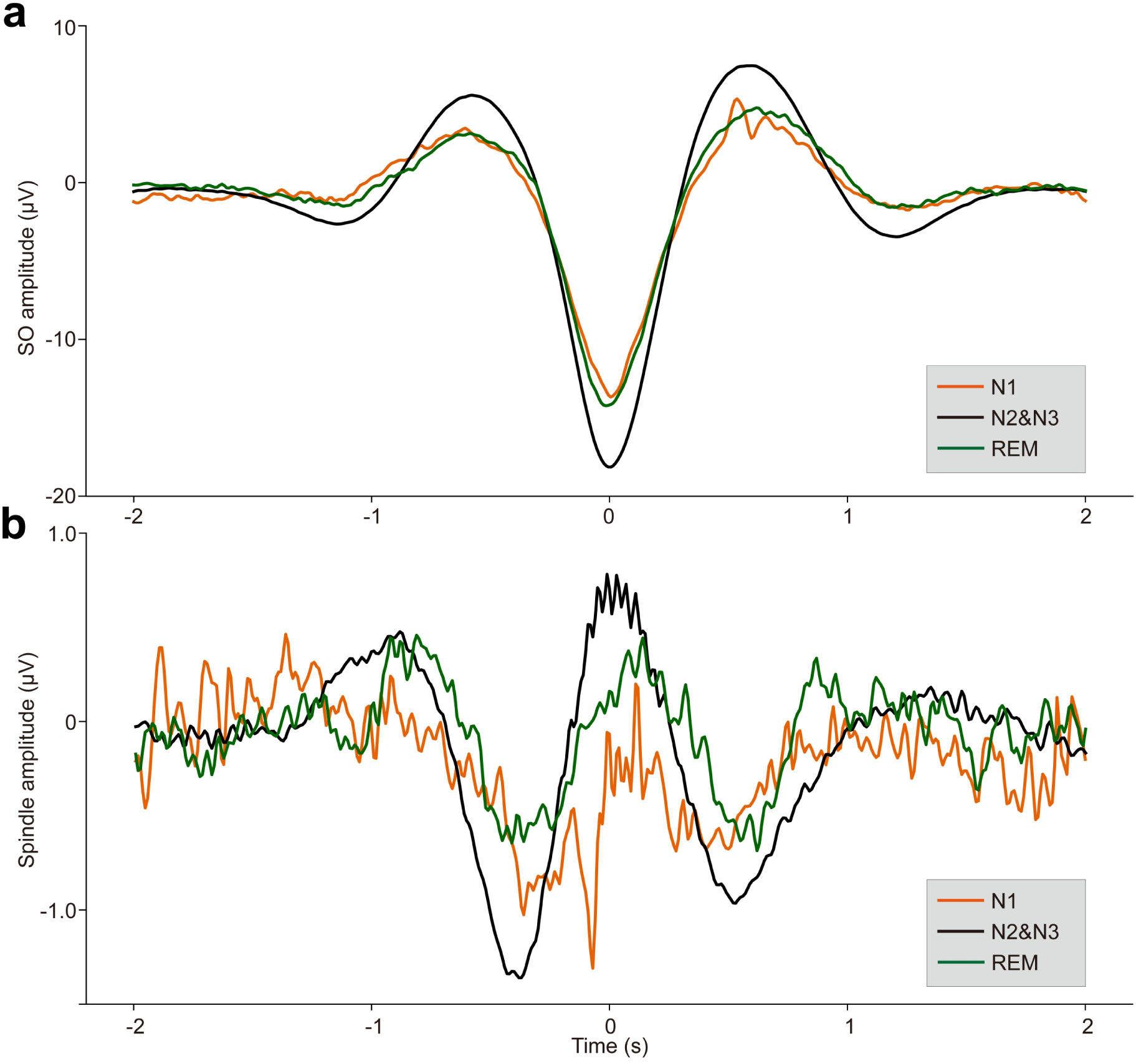
ERPs of SOs and spindles coupling during different sleep stages across all 107 subjects. **a.** ERP of SOs in different sleep stages using the broadband (0.1–30 Hz) EEG data. We align the trough of the DOWN-state of each SO at time zero (see Methods for details). The orange line represents the SO ERP in the N1 stage, the black line represents the SO ERP in the N2&N3 stage, and the green line represents the SO ERP in the REM stage. **b.** ERP of spindles in different sleep stages using the broadband (0.1–30 Hz) EEG data. We align the peak of each spindle at time zero (see Methods for details). The colour scheme is the same as in panel a.

**Fig. S3.**
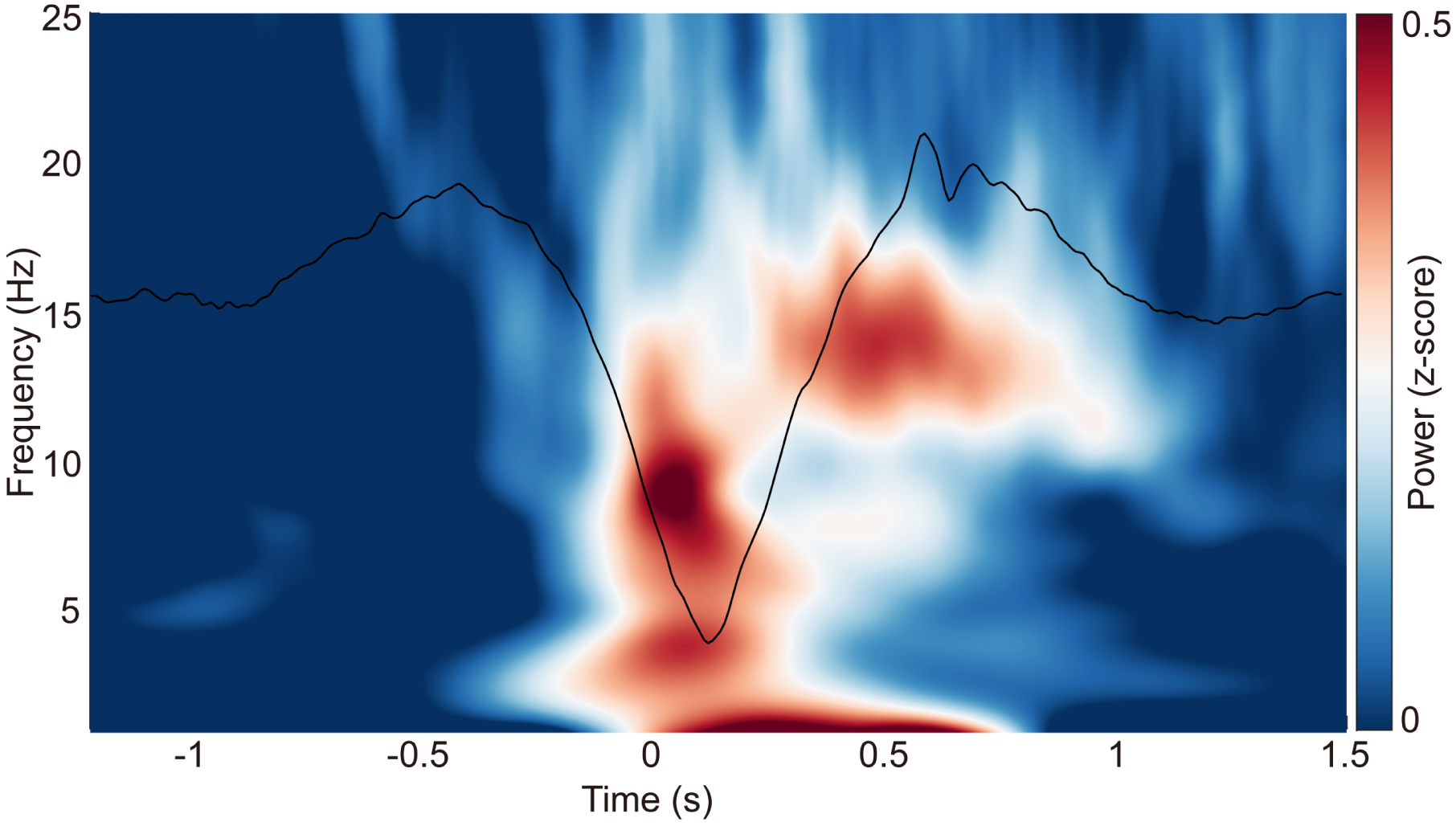
ERP and time-frequency patterns of SO-spindle coupling in the N1 stage. The averaged temporal frequency pattern and ERP across all instances of SO-spindle coupling, computed over all subjects, following the same procedure as in Fig. 2a, but for N1 stage.

**Fig. S4.**
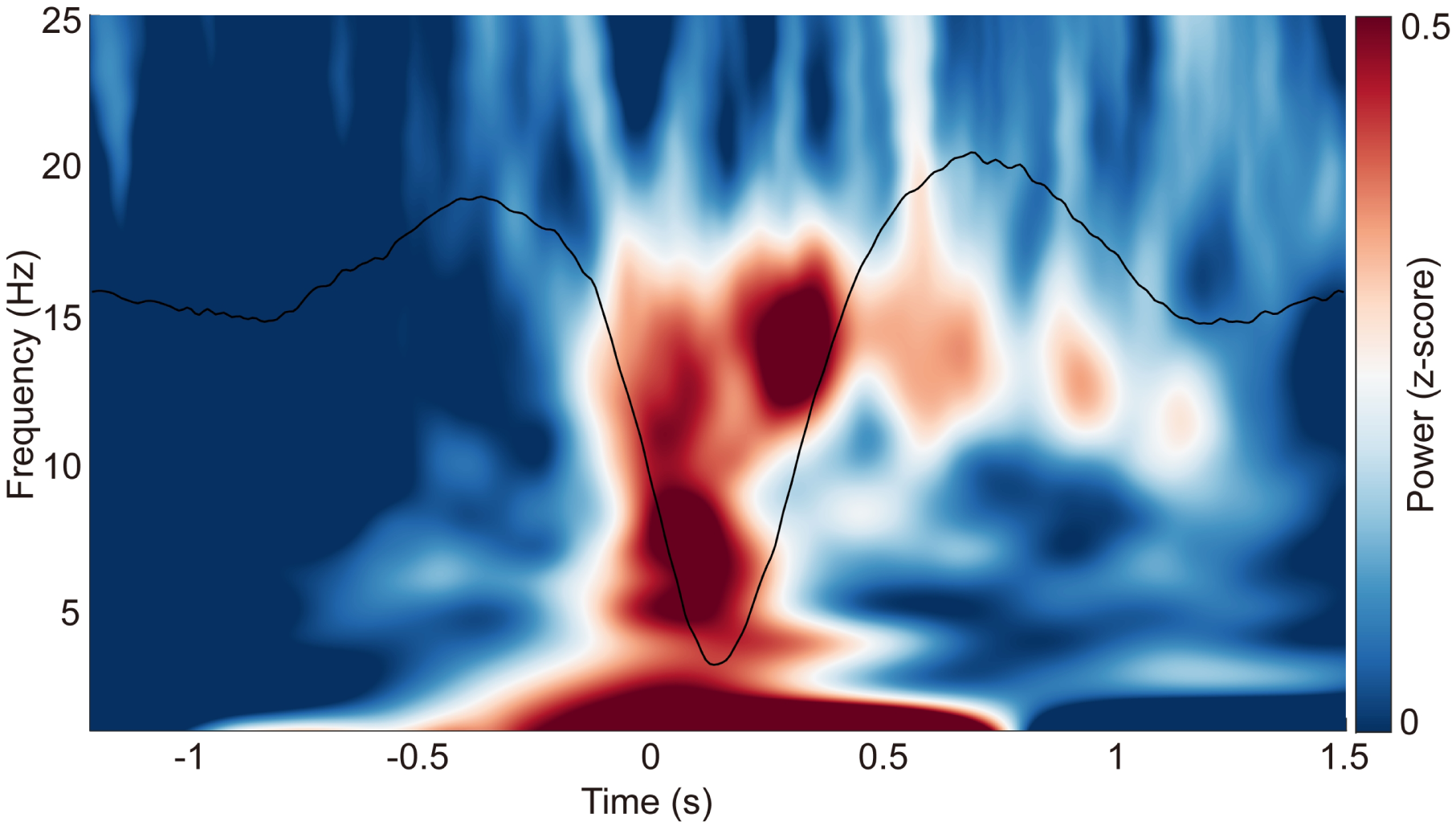
ERP and time-frequency patterns of SO-spindle coupling in the REM stage. The averaged temporal frequency pattern and ERP across all instances of SO-spindle coupling, computed over all subjects, again following the same procedure as in Fig. 2a, but for REM stage.

**Fig. S5.**
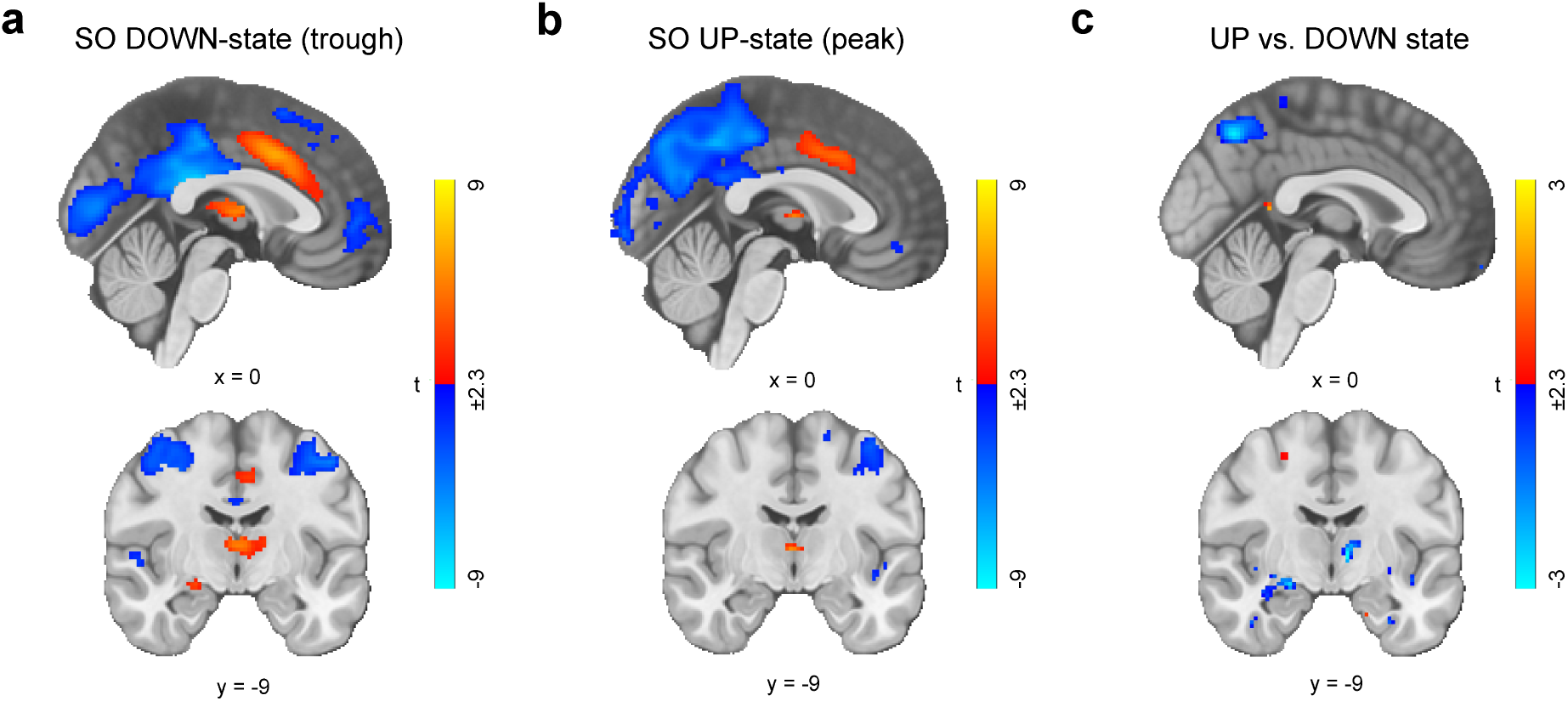
Brain-wide activity between SO UP-state (peak) and DOWN-state (trough). **a**, Brain activity with SO DOWN-state (trough) modelled as event onset, whole-brain FWE corrected at the cluster level (*p* < 0.05) with a cluster-forming voxel threshold of *p_unc._* < 0.001. **b**, Brain activity with SO UP-state (peak) modelled as event onset. **c**, Differences in brain activity corresponding to SO UP-state and SO DOWN-state. The whole-brain results were displayed at an uncorrected threshold of *p* < 0.01 for visualization purpose only. No brain region was found significant in this contrast.

**Fig. S6.**
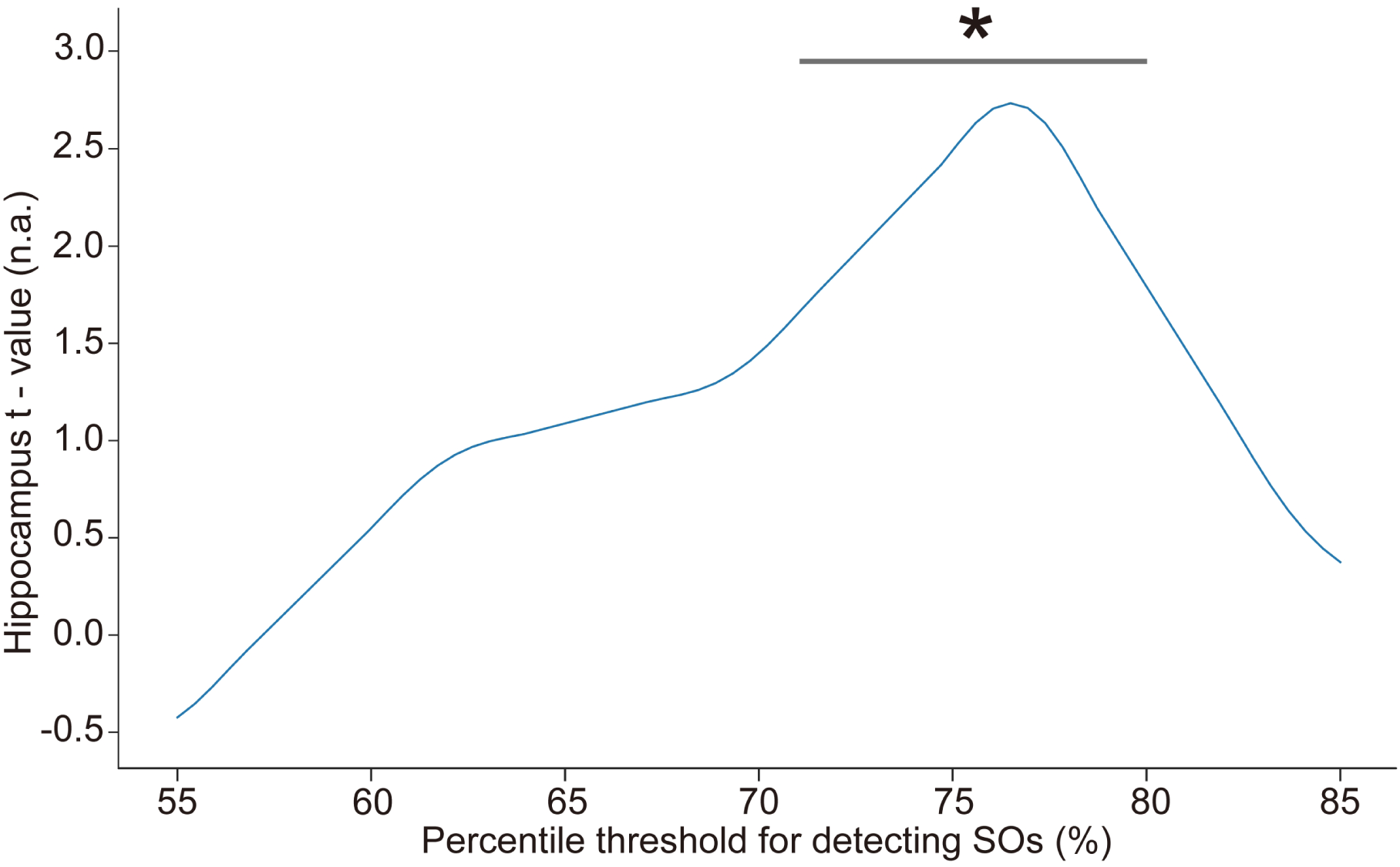
Influence of the percentile threshold for SO detection on hippocampal activation (ROI) during SO-spindle coupling. We changed the percentile threshold for SO event detection in the EEG data analysis and then reconstructed the GLM design matrix based on the SO events detected at each threshold. The brain-wide activation pattern of SO-spindle couplings in the N2/3 stage was extracted using the same method as shown in Fig. 3. The gray horizontal line represents the significant range (71%–80%). * *p* < 0.05.

**Fig. S7.**
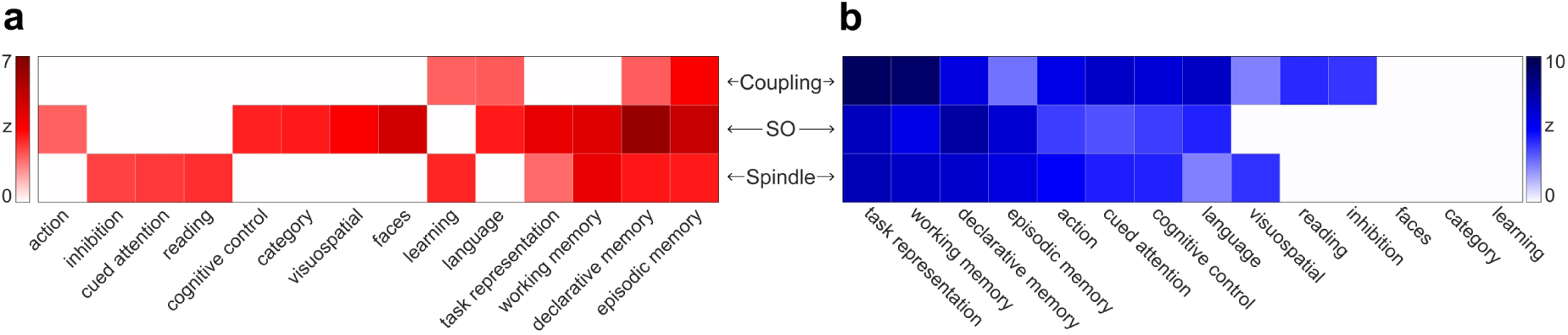
Functional decoding using the ROI association method in Neurosynth. **a**, Decoding results using positive activation. **b**, Decoding results using negative activation. Each row corresponds to the brain-wide activation patterns for sleep rhythms shown in Fig. 3b**-d**, while each column corresponds to topics in the Neurosynth database (detailed in Methods). Only topics with a decoded significance level of *p* < 0.05 are displayed.

**Table S1.**
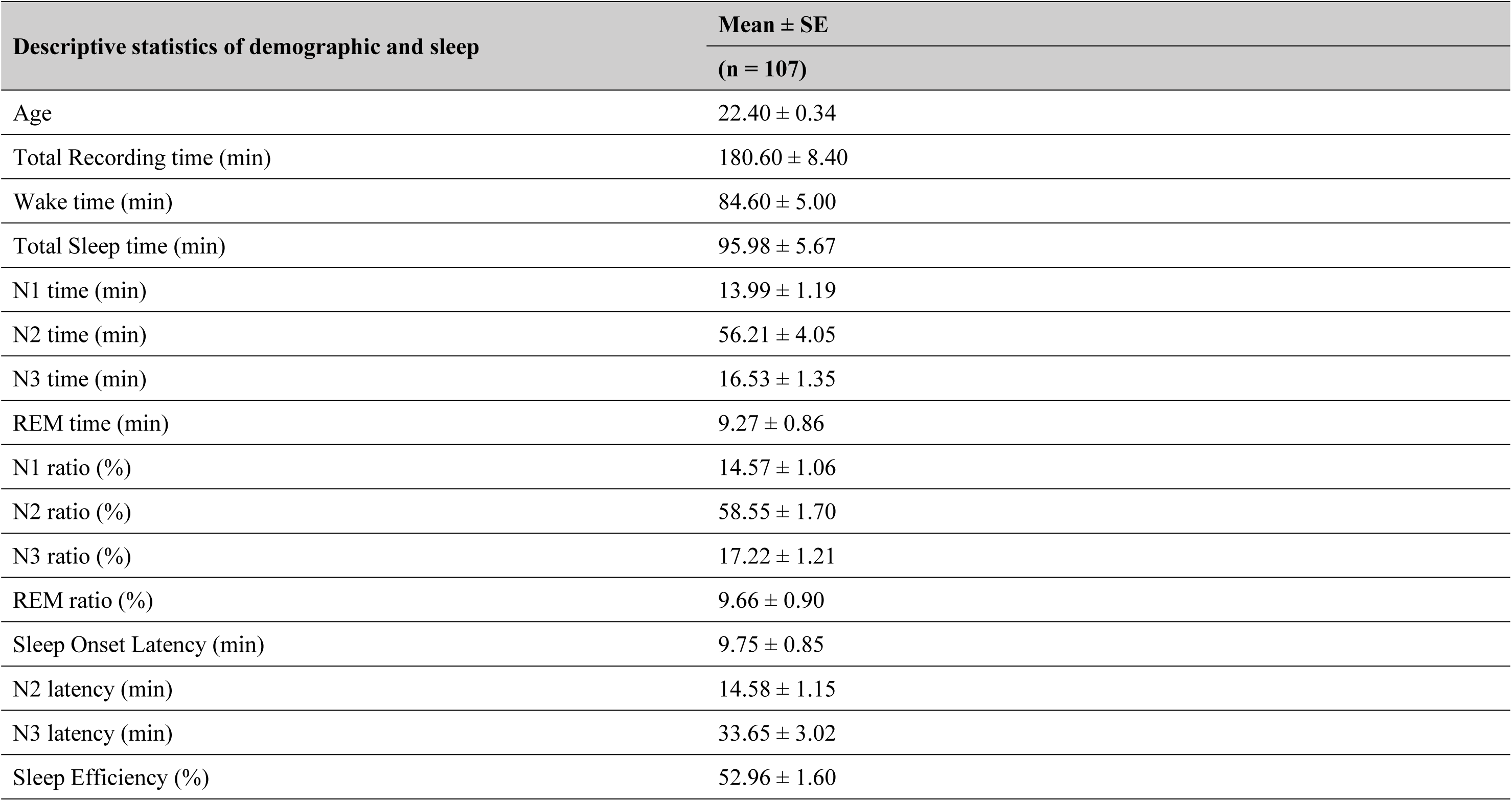
Descriptive results of demographic information and sleep characteristics. . Note: The total recorded time is equal to the awake time plus the total sleep time. The sleep onset latency is the time taken to reach the first sleep epoch. The Sleep Efficiency is the ratio of actual sleep time to total recording time.

| Descriptive statistics of demographic and sleep | Mean ± SE |
| --- | --- |
|  | (n = 107) |
| Age | 22.40 ± 0.34 |
| Total Recording time (min) | 180.60 ± 8.40 |
| Wake time (min) | 84.60 ± 5.00 |
| Total Sleep time (min) | 95.98 ± 5.67 |
| N1 time (min) | 13.99 ± 1.19 |
| N2 time (min) | 56.21 ± 4.05 |
| N3 time (min) | 16.53 ± 1.35 |
| REM time (min) | 9.27 ± 0.86 |
| N1 ratio (%) | 14.57 ± 1.06 |
| N2 ratio (%) | 58.55 ± 1.70 |
| N3 ratio (%) | 17.22 ± 1.21 |
| REM ratio (%) | 9.66 ± 0.90 |
| Sleep Onset Latency (min) | 9.75 ± 0.85 |
| N2 latency (min) | 14.58 ± 1.15 |
| N3 latency (min) | 33.65 ± 3.02 |
| Sleep Efficiency (%) | 52.96 ± 1.60 |

**Table S2.** Statistics of sleep duration, SO, spindle and coupling event numbers and densities for all 107 subjects during N1 stage.

| Subject ID | N1 Sleep Stage |  |  |  |  |  |  |
| --- | --- | --- | --- | --- | --- | --- | --- |
|  | Time (min) | Number of SO | Number of spindle | Number of coupling | Density of SO (times/min) | Density of spindle (times/min) | Density of coupling (times/min) |
| 1 | 8.38 | 27 | 24 | 6 | 3.22 | 2.87 | 0.72 |
| 2 | 5.21 | 26 | 23 | 8 | 4.99 | 4.41 | 1.54 |
| 3 | 3.24 | 14 | 14 | 4 | 4.32 | 4.32 | 1.23 |
| 4 | 16.61 | 25 | 49 | 6 | 1.51 | 2.95 | 0.36 |
| 5 | 3.38 | 0 | 0 | 0 | 0.00 | 0.00 | 0.00 |
| 6 | 9.74 | 35 | 34 | 11 | 3.59 | 3.49 | 1.13 |
| 7 | 8.84 | 25 | 16 | 11 | 2.83 | 1.81 | 1.24 |
| 8 | 19.51 | 96 | 116 | 40 | 4.92 | 5.95 | 2.05 |
| 9 | 2.28 | 5 | 2 | 1 | 2.20 | 0.88 | 0.44 |
| 10 | 25.71 | 29 | 11 | 3 | 1.13 | 0.43 | 0.12 |
| 11 | 8.04 | 24 | 19 | 5 | 2.98 | 2.36 | 0.62 |
| 12 | 4.54 | 6 | 5 | 1 | 1.32 | 1.10 | 0.22 |
| 13 | 4.91 | 15 | 22 | 5 | 3.05 | 4.48 | 1.02 |
| 14 | 3.41 | 3 | 15 | 1 | 0.88 | 4.40 | 0.29 |
| 15 | 4.91 | 10 | 9 | 2 | 2.04 | 1.83 | 0.41 |
| 16 | 21.98 | 39 | 113 | 14 | 1.77 | 5.14 | 0.64 |
| 17 | 1.31 | 8 | 7 | 4 | 6.11 | 5.34 | 3.05 |
| 18 | 0.91 | 0 | 0 | 0 | 0.00 | 0.00 | 0.00 |
| 19 | 2.54 | 10 | 8 | 4 | 3.93 | 3.15 | 1.57 |
| 20 | 23.71 | 78 | 64 | 23 | 3.29 | 2.70 | 0.97 |
| 21 | 35.81 | 72 | 50 | 13 | 2.01 | 1.40 | 0.36 |
| 22 | 25.58 | 42 | 103 | 20 | 1.64 | 4.03 | 0.78 |
| 23 | 3.41 | 0 | 0 | 0 | 0.00 | 0.00 | 0.00 |
| 24 | 20.68 | 48 | 50 | 13 | 2.32 | 2.42 | 0.63 |
| 25 | 47.74 | 142 | 192 | 51 | 2.97 | 4.02 | 1.07 |
| 26 | 20.91 | 66 | 43 | 16 | 3.16 | 2.06 | 0.77 |
| 27 | 7.91 | 48 | 38 | 12 | 6.07 | 4.80 | 1.52 |
| 28 | 10.71 | 30 | 42 | 12 | 2.80 | 3.92 | 1.12 |
| 29 | 22.71 | 99 | 74 | 31 | 4.36 | 3.26 | 1.37 |
| 30 | 2.84 | 14 | 12 | 2 | 4.92 | 4.22 | 0.70 |
| 31 | 7.41 | 17 | 6 | 2 | 2.29 | 0.81 | 0.27 |
| 32 | 0.98 | 0 | 0 | 0 | 0.00 | 0.00 | 0.00 |
| 33 | 3.21 | 20 | 12 | 4 | 6.23 | 3.74 | 1.25 |
| 34 | 7.81 | 32 | 13 | 4 | 4.10 | 1.66 | 0.51 |
| 35 | 4.44 | 14 | 6 | 2 | 3.15 | 1.35 | 0.45 |
| 36 | 41.34 | 123 | 101 | 28 | 2.98 | 2.44 | 0.68 |
| 37 | 5.31 | 27 | 17 | 12 | 5.08 | 3.20 | 2.26 |
| 38 | 13.34 | 39 | 38 | 13 | 2.92 | 2.85 | 0.97 |
| 39 | 20.34 | 64 | 36 | 8 | 3.15 | 1.77 | 0.39 |
| 40 | 5.74 | 28 | 16 | 6 | 4.88 | 2.79 | 1.04 |
| 41 | 8.11 | 1 | 16 | 1 | 0.12 | 1.97 | 0.12 |
| 42 | 25.68 | 63 | 105 | 17 | 2.45 | 4.09 | 0.66 |
| 43 | 23.21 | 46 | 17 | 9 | 1.98 | 0.73 | 0.39 |
| 44 | 16.94 | 44 | 66 | 10 | 2.60 | 3.90 | 0.59 |
| 45 | 6.31 | 11 | 20 | 3 | 1.74 | 3.17 | 0.48 |
| 46 | 7.24 | 41 | 20 | 5 | 5.66 | 2.76 | 0.69 |
| 47 | 9.14 | 32 | 27 | 11 | 3.50 | 2.95 | 1.20 |
| 48 | 35.71 | 139 | 177 | 54 | 3.89 | 4.96 | 1.51 |
| 49 | 7.01 | 15 | 27 | 7 | 2.14 | 3.85 | 1.00 |
| 50 | 27.18 | 117 | 55 | 18 | 4.31 | 2.02 | 0.66 |
| 51 | 13.38 | 42 | 53 | 13 | 3.14 | 3.96 | 0.97 |
| 52 | 17.54 | 48 | 42 | 4 | 2.74 | 2.39 | 0.23 |
| 53 | 24.31 | 140 | 78 | 30 | 5.76 | 3.21 | 1.23 |
| 54 | 11.54 | 37 | 60 | 13 | 3.21 | 5.20 | 1.13 |
| 55 | 10.14 | 2 | 27 | 1 | 0.20 | 2.66 | 0.10 |
| 56 | 4.41 | 27 | 21 | 9 | 6.12 | 4.76 | 2.04 |
| 57 | 5.01 | 19 | 3 | 1 | 3.79 | 0.60 | 0.20 |
| 58 | 6.01 | 28 | 28 | 7 | 4.66 | 4.66 | 1.16 |
| 59 | 37.38 | 136 | 93 | 30 | 3.64 | 2.49 | 0.80 |
| 60 | 7.08 | 35 | 9 | 3 | 4.95 | 1.27 | 0.42 |
| 61 | 8.74 | 38 | 36 | 9 | 4.35 | 4.12 | 1.03 |
| 62 | 17.61 | 45 | 24 | 12 | 2.56 | 1.36 | 0.68 |
| 63 | 48.61 | 125 | 148 | 31 | 2.57 | 3.04 | 0.64 |
| 64 | 9.74 | 29 | 54 | 10 | 2.98 | 5.54 | 1.03 |
| 65 | 22.21 | 40 | 47 | 9 | 1.80 | 2.12 | 0.41 |
| 66 | 36.78 | 63 | 52 | 14 | 1.71 | 1.41 | 0.38 |
| 67 | 33.51 | 37 | 54 | 13 | 1.10 | 1.61 | 0.39 |
| 68 | 10.71 | 50 | 11 | 3 | 4.67 | 1.03 | 0.28 |
| 69 | 58.34 | 143 | 130 | 31 | 2.45 | 2.23 | 0.53 |
| 70 | 14.21 | 50 | 55 | 16 | 3.52 | 3.87 | 1.13 |
| 71 | 21.98 | 88 | 74 | 26 | 4.00 | 3.37 | 1.18 |
| 72 | 55.31 | 212 | 203 | 45 | 3.83 | 3.67 | 0.81 |
| 73 | 24.61 | 51 | 19 | 5 | 2.07 | 0.77 | 0.20 |
| 74 | 23.31 | 100 | 47 | 27 | 4.29 | 2.02 | 1.16 |
| 75 | 4.74 | 10 | 7 | 2 | 2.11 | 1.48 | 0.42 |
| 76 | 37.11 | 77 | 34 | 9 | 2.07 | 0.92 | 0.24 |
| 77 | 14.41 | 55 | 25 | 12 | 3.82 | 1.73 | 0.83 |
| 78 | 12.21 | 0 | 0 | 0 | 0.00 | 0.00 | 0.00 |
| 79 | 6.18 | 0 | 0 | 0 | 0.00 | 0.00 | 0.00 |
| 80 | 8.18 | 2 | 16 | 1 | 0.24 | 1.96 | 0.12 |
| 81 | 2.84 | 9 | 5 | 2 | 3.17 | 1.76 | 0.70 |
| 82 | 7.41 | 19 | 21 | 6 | 2.56 | 2.83 | 0.81 |
| 83 | 1.91 | 11 | 5 | 2 | 5.76 | 2.62 | 1.05 |
| 84 | 2.14 | 11 | 4 | 1 | 5.13 | 1.87 | 0.47 |
| 85 | 10.38 | 24 | 38 | 7 | 2.31 | 3.66 | 0.67 |
| 86 | 10.34 | 43 | 55 | 14 | 4.16 | 5.32 | 1.35 |
| 87 | 5.98 | 20 | 16 | 3 | 3.35 | 2.68 | 0.50 |
| 88 | 0.41 | 0 | 0 | 0 | 0.00 | 0.00 | 0.00 |
| 89 | 10.91 | 16 | 48 | 5 | 1.47 | 4.40 | 0.46 |
| 90 | 9.54 | 45 | 28 | 11 | 4.72 | 2.93 | 1.15 |
| 91 | 4.18 | 26 | 11 | 3 | 6.23 | 2.63 | 0.72 |
| 92 | 2.14 | 0 | 0 | 0 | 0.00 | 0.00 | 0.00 |
| 93 | 10.11 | 26 | 35 | 9 | 2.57 | 3.46 | 0.89 |
| 94 | 1.91 | 3 | 10 | 2 | 1.57 | 5.24 | 1.05 |
| 95 | 8.68 | 11 | 24 | 4 | 1.27 | 2.77 | 0.46 |
| 96 | 30.71 | 112 | 99 | 28 | 3.65 | 3.22 | 0.91 |
| 97 | 11.71 | 22 | 15 | 5 | 1.88 | 1.28 | 0.43 |
| 98 | 11.91 | 42 | 31 | 8 | 3.53 | 2.60 | 0.67 |
| 99 | 16.41 | 42 | 69 | 15 | 2.56 | 4.20 | 0.91 |
| 100 | 7.34 | 23 | 27 | 5 | 3.13 | 3.68 | 0.68 |
| 101 | 6.41 | 13 | 18 | 3 | 2.03 | 2.81 | 0.47 |
| 102 | 9.24 | 33 | 28 | 7 | 3.57 | 3.03 | 0.76 |
| 103 | 3.34 | 20 | 12 | 5 | 5.98 | 3.59 | 1.50 |
| 104 | 13.41 | 30 | 66 | 9 | 2.24 | 4.92 | 0.67 |
| 105 | 10.58 | 5 | 26 | 3 | 0.47 | 2.46 | 0.28 |
| 106 | 6.48 | 22 | 20 | 9 | 3.40 | 3.09 | 1.39 |
| 107 | 21.51 | 65 | 6 | 2 | 3.02 | 0.28 | 0.09 |
| **Mean** | 13.99 | 40.48 | 38.29 | 10.16 | 2.95 | 2.71 | 0.75 |
| **SE** | 1.19 | 3.85 | 3.92 | 1.06 | 0.16 | 0.14 | 0.05 |

**Table S3.**
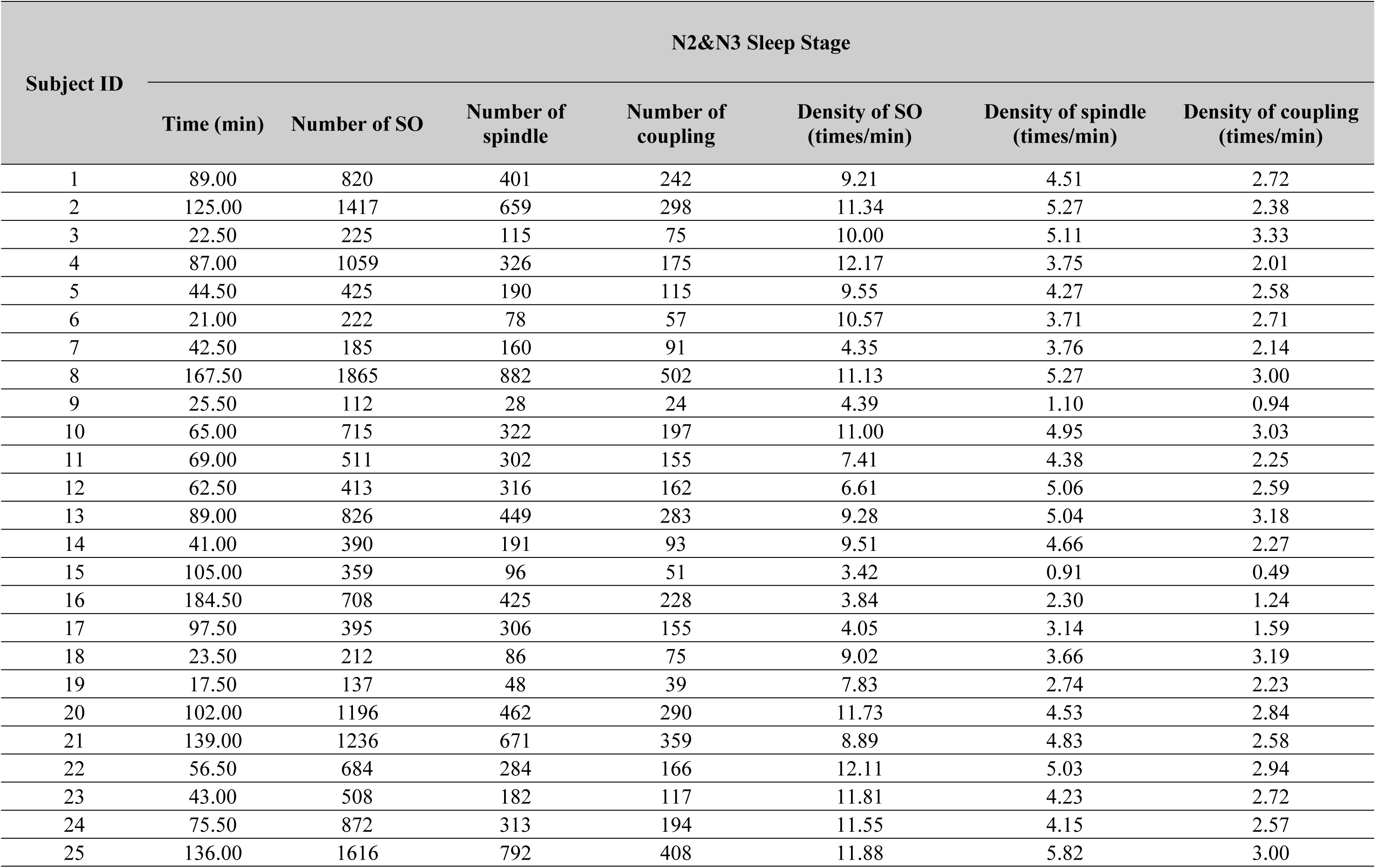
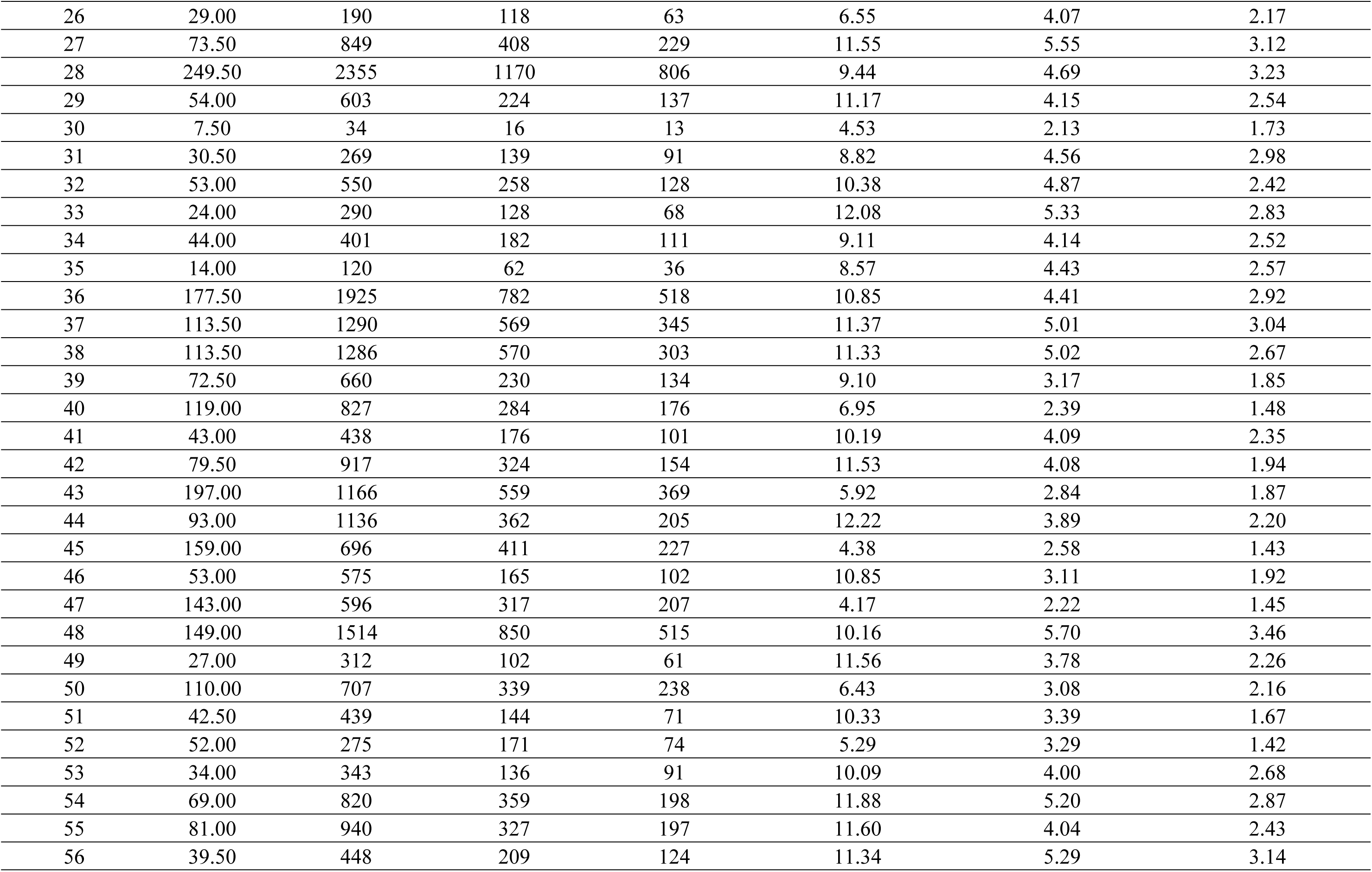
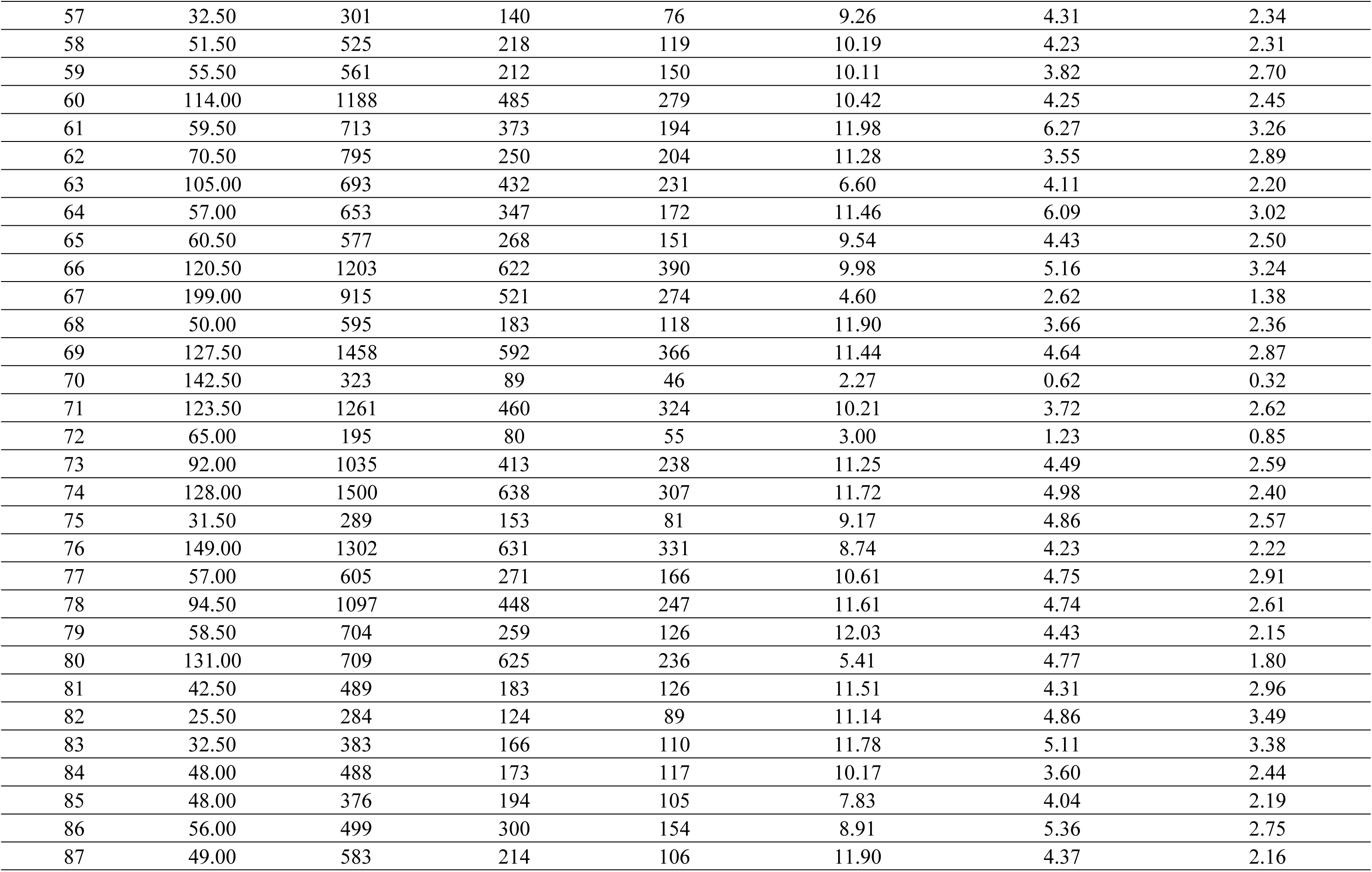
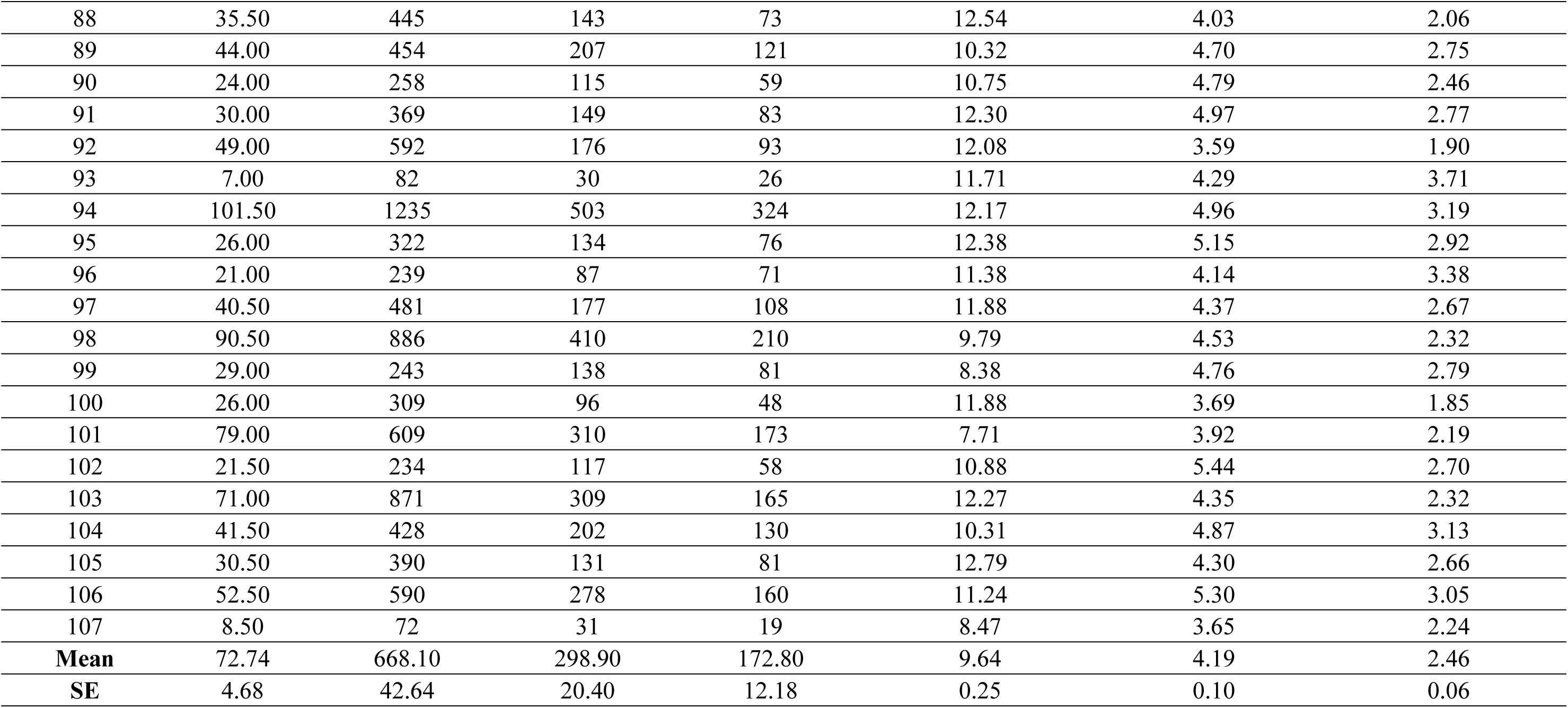
Statistics of sleep duration, SO, spindle and coupling event numbers and densities for all 107 subjects during N2&N3 stage.

| Subject ID | N2&N3 Sleep Stage |  |  |  |  |  |  |
| --- | --- | --- | --- | --- | --- | --- | --- |
|  | Time (min) | Number of SO | Number of spindle | Number of coupling | Density of SO (times/min) | Density of spindle (times/min) | Density of coupling (times/min) |
| 1 | 89.00 | 820 | 401 | 242 | 9.21 | 4.51 | 2.72 |
| 2 | 125.00 | 1417 | 659 | 298 | 11.34 | 5.27 | 2.38 |
| 3 | 22.50 | 225 | 115 | 75 | 10.00 | 5.11 | 3.33 |
| 4 | 87.00 | 1059 | 326 | 175 | 12.17 | 3.75 | 2.01 |
| 5 | 44.50 | 425 | 190 | 115 | 9.55 | 4.27 | 2.58 |
| 6 | 21.00 | 222 | 78 | 57 | 10.57 | 3.71 | 2.71 |
| 7 | 42.50 | 185 | 160 | 91 | 4.35 | 3.76 | 2.14 |
| 8 | 167.50 | 1865 | 882 | 502 | 11.13 | 5.27 | 3.00 |
| 9 | 25.50 | 112 | 28 | 24 | 4.39 | 1.10 | 0.94 |
| 10 | 65.00 | 715 | 322 | 197 | 11.00 | 4.95 | 3.03 |
| 11 | 69.00 | 511 | 302 | 155 | 7.41 | 4.38 | 2.25 |
| 12 | 62.50 | 413 | 316 | 162 | 6.61 | 5.06 | 2.59 |
| 13 | 89.00 | 826 | 449 | 283 | 9.28 | 5.04 | 3.18 |
| 14 | 41.00 | 390 | 191 | 93 | 9.51 | 4.66 | 2.27 |
| 15 | 105.00 | 359 | 96 | 51 | 3.42 | 0.91 | 0.49 |
| 16 | 184.50 | 708 | 425 | 228 | 3.84 | 2.30 | 1.24 |
| 17 | 97.50 | 395 | 306 | 155 | 4.05 | 3.14 | 1.59 |
| 18 | 23.50 | 212 | 86 | 75 | 9.02 | 3.66 | 3.19 |
| 19 | 17.50 | 137 | 48 | 39 | 7.83 | 2.74 | 2.23 |
| 20 | 102.00 | 1196 | 462 | 290 | 11.73 | 4.53 | 2.84 |
| 21 | 139.00 | 1236 | 671 | 359 | 8.89 | 4.83 | 2.58 |
| 22 | 56.50 | 684 | 284 | 166 | 12.11 | 5.03 | 2.94 |
| 23 | 43.00 | 508 | 182 | 117 | 11.81 | 4.23 | 2.72 |
| 24 | 75.50 | 872 | 313 | 194 | 11.55 | 4.15 | 2.57 |
| 25 | 136.00 | 1616 | 792 | 408 | 11.88 | 5.82 | 3.00 |
| 26 | 29.00 | 190 | 118 | 63 | 6.55 | 4.07 | 2.17 |
| 27 | 73.50 | 849 | 408 | 229 | 11.55 | 5.55 | 3.12 |
| 28 | 249.50 | 2355 | 1170 | 806 | 9.44 | 4.69 | 3.23 |
| 29 | 54.00 | 603 | 224 | 137 | 11.17 | 4.15 | 2.54 |
| 30 | 7.50 | 34 | 16 | 13 | 4.53 | 2.13 | 1.73 |
| 31 | 30.50 | 269 | 139 | 91 | 8.82 | 4.56 | 2.98 |
| 32 | 53.00 | 550 | 258 | 128 | 10.38 | 4.87 | 2.42 |
| 33 | 24.00 | 290 | 128 | 68 | 12.08 | 5.33 | 2.83 |
| 34 | 44.00 | 401 | 182 | 111 | 9.11 | 4.14 | 2.52 |
| 35 | 14.00 | 120 | 62 | 36 | 8.57 | 4.43 | 2.57 |
| 36 | 177.50 | 1925 | 782 | 518 | 10.85 | 4.41 | 2.92 |
| 37 | 113.50 | 1290 | 569 | 345 | 11.37 | 5.01 | 3.04 |
| 38 | 113.50 | 1286 | 570 | 303 | 11.33 | 5.02 | 2.67 |
| 39 | 72.50 | 660 | 230 | 134 | 9.10 | 3.17 | 1.85 |
| 40 | 119.00 | 827 | 284 | 176 | 6.95 | 2.39 | 1.48 |
| 41 | 43.00 | 438 | 176 | 101 | 10.19 | 4.09 | 2.35 |
| 42 | 79.50 | 917 | 324 | 154 | 11.53 | 4.08 | 1.94 |
| 43 | 197.00 | 1166 | 559 | 369 | 5.92 | 2.84 | 1.87 |
| 44 | 93.00 | 1136 | 362 | 205 | 12.22 | 3.89 | 2.20 |
| 45 | 159.00 | 696 | 411 | 227 | 4.38 | 2.58 | 1.43 |
| 46 | 53.00 | 575 | 165 | 102 | 10.85 | 3.11 | 1.92 |
| 47 | 143.00 | 596 | 317 | 207 | 4.17 | 2.22 | 1.45 |
| 48 | 149.00 | 1514 | 850 | 515 | 10.16 | 5.70 | 3.46 |
| 49 | 27.00 | 312 | 102 | 61 | 11.56 | 3.78 | 2.26 |
| 50 | 110.00 | 707 | 339 | 238 | 6.43 | 3.08 | 2.16 |
| 51 | 42.50 | 439 | 144 | 71 | 10.33 | 3.39 | 1.67 |
| 52 | 52.00 | 275 | 171 | 74 | 5.29 | 3.29 | 1.42 |
| 53 | 34.00 | 343 | 136 | 91 | 10.09 | 4.00 | 2.68 |
| 54 | 69.00 | 820 | 359 | 198 | 11.88 | 5.20 | 2.87 |
| 55 | 81.00 | 940 | 327 | 197 | 11.60 | 4.04 | 2.43 |
| 56 | 39.50 | 448 | 209 | 124 | 11.34 | 5.29 | 3.14 |
| 57 | 32.50 | 301 | 140 | 76 | 9.26 | 4.31 | 2.34 |
| 58 | 51.50 | 525 | 218 | 119 | 10.19 | 4.23 | 2.31 |
| 59 | 55.50 | 561 | 212 | 150 | 10.11 | 3.82 | 2.70 |
| 60 | 114.00 | 1188 | 485 | 279 | 10.42 | 4.25 | 2.45 |
| 61 | 59.50 | 713 | 373 | 194 | 11.98 | 6.27 | 3.26 |
| 62 | 70.50 | 795 | 250 | 204 | 11.28 | 3.55 | 2.89 |
| 63 | 105.00 | 693 | 432 | 231 | 6.60 | 4.11 | 2.20 |
| 64 | 57.00 | 653 | 347 | 172 | 11.46 | 6.09 | 3.02 |
| 65 | 60.50 | 577 | 268 | 151 | 9.54 | 4.43 | 2.50 |
| 66 | 120.50 | 1203 | 622 | 390 | 9.98 | 5.16 | 3.24 |
| 67 | 199.00 | 915 | 521 | 274 | 4.60 | 2.62 | 1.38 |
| 68 | 50.00 | 595 | 183 | 118 | 11.90 | 3.66 | 2.36 |
| 69 | 127.50 | 1458 | 592 | 366 | 11.44 | 4.64 | 2.87 |
| 70 | 142.50 | 323 | 89 | 46 | 2.27 | 0.62 | 0.32 |
| 71 | 123.50 | 1261 | 460 | 324 | 10.21 | 3.72 | 2.62 |
| 72 | 65.00 | 195 | 80 | 55 | 3.00 | 1.23 | 0.85 |
| 73 | 92.00 | 1035 | 413 | 238 | 11.25 | 4.49 | 2.59 |
| 74 | 128.00 | 1500 | 638 | 307 | 11.72 | 4.98 | 2.40 |
| 75 | 31.50 | 289 | 153 | 81 | 9.17 | 4.86 | 2.57 |
| 76 | 149.00 | 1302 | 631 | 331 | 8.74 | 4.23 | 2.22 |
| 77 | 57.00 | 605 | 271 | 166 | 10.61 | 4.75 | 2.91 |
| 78 | 94.50 | 1097 | 448 | 247 | 11.61 | 4.74 | 2.61 |
| 79 | 58.50 | 704 | 259 | 126 | 12.03 | 4.43 | 2.15 |
| 80 | 131.00 | 709 | 625 | 236 | 5.41 | 4.77 | 1.80 |
| 81 | 42.50 | 489 | 183 | 126 | 11.51 | 4.31 | 2.96 |
| 82 | 25.50 | 284 | 124 | 89 | 11.14 | 4.86 | 3.49 |
| 83 | 32.50 | 383 | 166 | 110 | 11.78 | 5.11 | 3.38 |
| 84 | 48.00 | 488 | 173 | 117 | 10.17 | 3.60 | 2.44 |
| 85 | 48.00 | 376 | 194 | 105 | 7.83 | 4.04 | 2.19 |
| 86 | 56.00 | 499 | 300 | 154 | 8.91 | 5.36 | 2.75 |
| 87 | 49.00 | 583 | 214 | 106 | 11.90 | 4.37 | 2.16 |
| 88 | 35.50 | 445 | 143 | 73 | 12.54 | 4.03 | 2.06 |
| 89 | 44.00 | 454 | 207 | 121 | 10.32 | 4.70 | 2.75 |
| 90 | 24.00 | 258 | 115 | 59 | 10.75 | 4.79 | 2.46 |
| 91 | 30.00 | 369 | 149 | 83 | 12.30 | 4.97 | 2.77 |
| 92 | 49.00 | 592 | 176 | 93 | 12.08 | 3.59 | 1.90 |
| 93 | 7.00 | 82 | 30 | 26 | 11.71 | 4.29 | 3.71 |
| 94 | 101.50 | 1235 | 503 | 324 | 12.17 | 4.96 | 3.19 |
| 95 | 26.00 | 322 | 134 | 76 | 12.38 | 5.15 | 2.92 |
| 96 | 21.00 | 239 | 87 | 71 | 11.38 | 4.14 | 3.38 |
| 97 | 40.50 | 481 | 177 | 108 | 11.88 | 4.37 | 2.67 |
| 98 | 90.50 | 886 | 410 | 210 | 9.79 | 4.53 | 2.32 |
| 99 | 29.00 | 243 | 138 | 81 | 8.38 | 4.76 | 2.79 |
| 100 | 26.00 | 309 | 96 | 48 | 11.88 | 3.69 | 1.85 |
| 101 | 79.00 | 609 | 310 | 173 | 7.71 | 3.92 | 2.19 |
| 102 | 21.50 | 234 | 117 | 58 | 10.88 | 5.44 | 2.70 |
| 103 | 71.00 | 871 | 309 | 165 | 12.27 | 4.35 | 2.32 |
| 104 | 41.50 | 428 | 202 | 130 | 10.31 | 4.87 | 3.13 |
| 105 | 30.50 | 390 | 131 | 81 | 12.79 | 4.30 | 2.66 |
| 106 | 52.50 | 590 | 278 | 160 | 11.24 | 5.30 | 3.05 |
| 107 | 8.50 | 72 | 31 | 19 | 8.47 | 3.65 | 2.24 |
| **Mean** | 72.74 | 668.10 | 298.90 | 172.80 | 9.64 | 4.19 | 2.46 |
| **SE** | 4.68 | 42.64 | 20.40 | 12.18 | 0.25 | 0.10 | 0.06 |

**Table S4.** Statistics of sleep duration, SO, spindle and coupling event numbers and densities for all 107 subjects during REM stage.

| Subject ID | REM Sleep Stage |  |  |  |  |  |  |
| --- | --- | --- | --- | --- | --- | --- | --- |
|  | Time (min) | Number of SO | Number of spindle | Number of coupling | Density of SO (times/min) | Density of spindle (times/min) | Density of coupling (times/min) |
| 1 | 2.00 | 3 | 7 | 1 | 1.50 | 3.50 | 0.50 |
| 2 | 4.00 | 15 | 13 | 5 | 3.75 | 3.25 | 1.25 |
| 3 | 4.50 | 14 | 8 | 4 | 3.11 | 1.78 | 0.89 |
| 4 | 7.00 | 0 | 0 | 0 | 0.00 | 0.00 | 0.00 |
| 5 | 3.50 | 11 | 11 | 2 | 3.14 | 3.14 | 0.57 |
| 6 | 24.50 | 28 | 29 | 3 | 1.14 | 1.18 | 0.12 |
| 7 | 5.00 | 0 | 0 | 0 | 0.00 | 0.00 | 0.00 |
| 8 | 21.00 | 5 | 15 | 2 | 0.24 | 0.71 | 0.10 |
| 9 | 4.50 | 0 | 0 | 0 | 0.00 | 0.00 | 0.00 |
| 10 | 15.00 | 39 | 19 | 6 | 2.60 | 1.27 | 0.40 |
| 11 | 4.50 | 0 | 0 | 0 | 0.00 | 0.00 | 0.00 |
| 12 | 4.50 | 12 | 6 | 1 | 2.67 | 1.33 | 0.22 |
| 13 | 2.00 | 0 | 0 | 0 | 0.00 | 0.00 | 0.00 |
| 14 | 0.50 | 0 | 0 | 0 | 0.00 | 0.00 | 0.00 |
| 15 | 16.00 | 38 | 36 | 4 | 2.38 | 2.25 | 0.25 |
| 16 | 11.50 | 8 | 20 | 3 | 0.70 | 1.74 | 0.26 |
| 17 | 12.50 | 5 | 18 | 1 | 0.40 | 1.44 | 0.08 |
| 18 | 4.00 | 0 | 0 | 0 | 0.00 | 0.00 | 0.00 |
| 19 | 11.50 | 59 | 20 | 10 | 5.13 | 1.74 | 0.87 |
| 20 | 5.50 | 31 | 15 | 5 | 5.64 | 2.73 | 0.91 |
| 21 | 12.00 | 19 | 34 | 5 | 1.58 | 2.83 | 0.42 |
| 22 | 0.50 | 0 | 0 | 0 | 0.00 | 0.00 | 0.00 |
| 23 | 11.50 | 0 | 0 | 0 | 0.00 | 0.00 | 0.00 |
| 24 | 8.00 | 28 | 4 | 4 | 3.50 | 0.50 | 0.50 |
| 25 | 32.50 | 11 | 3 | 1 | 0.34 | 0.09 | 0.03 |
| 26 | 2.50 | 6 | 6 | 1 | 2.40 | 2.40 | 0.40 |
| 27 | 7.50 | 40 | 8 | 3 | 5.33 | 1.07 | 0.40 |
| 28 | 5.50 | 12 | 17 | 7 | 2.18 | 3.09 | 1.27 |
| 29 | 17.50 | 23 | 69 | 13 | 1.31 | 3.94 | 0.74 |
| 30 | 6.00 | 26 | 16 | 6 | 4.33 | 2.67 | 1.00 |
| 31 | 13.00 | 30 | 12 | 4 | 2.31 | 0.92 | 0.31 |
| 32 | 0.50 | 0 | 0 | 0 | 0.00 | 0.00 | 0.00 |
| 33 | 1.00 | 0 | 0 | 0 | 0.00 | 0.00 | 0.00 |
| 34 | 9.50 | 42 | 7 | 1 | 4.42 | 0.74 | 0.11 |
| 35 | 2.50 | 10 | 9 | 4 | 4.00 | 3.60 | 1.60 |
| 36 | 9.50 | 37 | 21 | 6 | 3.89 | 2.21 | 0.63 |
| 37 | 24.00 | 76 | 40 | 9 | 3.17 | 1.67 | 0.38 |
| 38 | 4.00 | 2 | 6 | 2 | 0.50 | 1.50 | 0.50 |
| 39 | 13.00 | 49 | 19 | 5 | 3.77 | 1.46 | 0.38 |
| 40 | 8.50 | 15 | 18 | 4 | 1.76 | 2.12 | 0.47 |
| 41 | 7.00 | 8 | 16 | 4 | 1.14 | 2.29 | 0.57 |
| 42 | 17.00 | 45 | 67 | 11 | 2.65 | 3.94 | 0.65 |
| 43 | 25.00 | 27 | 107 | 14 | 1.08 | 4.28 | 0.56 |
| 44 | 15.50 | 33 | 55 | 5 | 2.13 | 3.55 | 0.32 |
| 45 | 7.50 | 18 | 21 | 5 | 2.40 | 2.80 | 0.67 |
| 46 | 0.50 | 0 | 0 | 0 | 0.00 | 0.00 | 0.00 |
| 47 | 1.00 | 0 | 0 | 0 | 0.00 | 0.00 | 0.00 |
| 48 | 46.00 | 129 | 236 | 49 | 2.80 | 5.13 | 1.07 |
| 49 | 0.50 | 0 | 0 | 0 | 0.00 | 0.00 | 0.00 |
| 50 | 29.50 | 69 | 74 | 19 | 2.34 | 2.51 | 0.64 |
| 51 | 19.50 | 49 | 43 | 11 | 2.51 | 2.21 | 0.56 |
| 52 | 8.00 | 0 | 0 | 0 | 0.00 | 0.00 | 0.00 |
| 53 | 6.00 | 20 | 5 | 2 | 3.33 | 0.83 | 0.33 |
| 54 | 6.00 | 5 | 24 | 4 | 0.83 | 4.00 | 0.67 |
| 55 | 1.00 | 0 | 0 | 0 | 0.00 | 0.00 | 0.00 |
| 56 | 20.00 | 32 | 21 | 5 | 1.60 | 1.05 | 0.25 |
| 57 | 0.50 | 0 | 0 | 0 | 0.00 | 0.00 | 0.00 |
| 58 | 6.00 | 14 | 22 | 3 | 2.33 | 3.67 | 0.50 |
| 59 | 45.00 | 220 | 146 | 65 | 4.89 | 3.24 | 1.44 |
| 60 | 0.50 | 0 | 0 | 0 | 0.00 | 0.00 | 0.00 |
| 61 | 8.00 | 0 | 0 | 0 | 0.00 | 0.00 | 0.00 |
| 62 | 10.50 | 41 | 12 | 5 | 3.90 | 1.14 | 0.48 |
| 63 | 5.50 | 19 | 13 | 2 | 3.45 | 2.36 | 0.36 |
| 64 | 7.50 | 20 | 41 | 6 | 2.67 | 5.47 | 0.80 |
| 65 | 9.00 | 15 | 35 | 2 | 1.67 | 3.89 | 0.22 |
| 66 | 22.00 | 58 | 32 | 13 | 2.64 | 1.45 | 0.59 |
| 67 | 22.00 | 108 | 63 | 19 | 4.91 | 2.86 | 0.86 |
| 68 | 18.50 | 101 | 16 | 5 | 5.46 | 0.86 | 0.27 |
| 69 | 5.50 | 13 | 10 | 1 | 2.36 | 1.82 | 0.18 |
| 70 | 10.00 | 13 | 15 | 2 | 1.30 | 1.50 | 0.20 |
| 71 | 29.50 | 69 | 69 | 15 | 2.34 | 2.34 | 0.51 |
| 72 | 0.00 | 0 | 0 | 0 | 0.00 | 0.00 | 0.00 |
| 73 | 14.50 | 44 | 6 | 4 | 3.03 | 0.41 | 0.28 |
| 74 | 13.50 | 45 | 27 | 12 | 3.33 | 2.00 | 0.89 |
| 75 | 3.50 | 8 | 5 | 1 | 2.29 | 1.43 | 0.29 |
| 76 | 4.50 | 6 | 2 | 1 | 1.33 | 0.44 | 0.22 |
| 77 | 14.00 | 49 | 23 | 11 | 3.50 | 1.64 | 0.79 |
| 78 | 26.00 | 86 | 25 | 5 | 3.31 | 0.96 | 0.19 |
| 79 | 2.50 | 8 | 9 | 2 | 3.20 | 3.60 | 0.80 |
| 80 | 6.50 | 4 | 10 | 1 | 0.62 | 1.54 | 0.15 |
| 81 | 4.00 | 2 | 15 | 1 | 0.50 | 3.75 | 0.25 |
| 82 | 3.00 | 13 | 10 | 4 | 4.33 | 3.33 | 1.33 |
| 83 | 10.50 | 34 | 26 | 8 | 3.24 | 2.48 | 0.76 |
| 84 | 1.00 | 6 | 1 | 1 | 6.00 | 1.00 | 1.00 |
| 85 | 8.00 | 22 | 28 | 6 | 2.75 | 3.50 | 0.75 |
| 86 | 0.00 | 0 | 0 | 0 | 0.00 | 0.00 | 0.00 |
| 87 | 3.50 | 4 | 5 | 1 | 1.14 | 1.43 | 0.29 |
| 88 | 1.50 | 9 | 2 | 1 | 6.00 | 1.33 | 0.67 |
| 89 | 8.50 | 13 | 29 | 3 | 1.53 | 3.41 | 0.35 |
| 90 | 4.00 | 22 | 11 | 3 | 5.50 | 2.75 | 0.75 |
| 91 | 1.50 | 8 | 4 | 2 | 5.33 | 2.67 | 1.33 |
| 92 | 2.00 | 5 | 8 | 1 | 2.50 | 4.00 | 0.50 |
| 93 | 1.50 | 0 | 0 | 0 | 0.00 | 0.00 | 0.00 |
| 94 | 5.50 | 10 | 22 | 3 | 1.82 | 4.00 | 0.55 |
| 95 | 6.00 | 6 | 17 | 2 | 1.00 | 2.83 | 0.33 |
| 96 | 7.00 | 20 | 23 | 4 | 2.86 | 3.29 | 0.57 |
| 97 | 0.00 | 0 | 0 | 0 | 0.00 | 0.00 | 0.00 |
| 98 | 11.00 | 12 | 26 | 3 | 1.09 | 2.36 | 0.27 |
| 99 | 15.00 | 20 | 24 | 2 | 1.33 | 1.60 | 0.13 |
| 100 | 4.50 | 5 | 18 | 2 | 1.11 | 4.00 | 0.44 |
| 101 | 3.50 | 4 | 8 | 2 | 1.14 | 2.29 | 0.57 |
| 102 | 7.50 | 27 | 19 | 7 | 3.60 | 2.53 | 0.93 |
| 103 | 1.00 | 0 | 0 | 0 | 0.00 | 0.00 | 0.00 |
| 104 | 8.50 | 16 | 33 | 8 | 1.88 | 3.88 | 0.94 |
| 105 | 6.50 | 0 | 0 | 0 | 0.00 | 0.00 | 0.00 |
| 106 | 2.50 | 8 | 11 | 3 | 3.20 | 4.40 | 1.20 |
| 107 | 16.50 | 80 | 6 | 1 | 4.85 | 0.36 | 0.06 |
| **Mean** | 9.27 | 22.58 | 19.64 | 4.62 | 2.07 | 1.81 | 0.43 |
| **SE** | 0.87 | 3.08 | 2.99 | 0.81 | 0.17 | 0.14 | 0.04 |

**Table S5.** Peak and significant cluster of fMRI activity during SO main effect. We used the SO main effect whole-brain activation patterns in **Fig. 3b**. ROIs were defined anatomically (see Methods). Cluster sizes are reported *p_unc._* < 0.001.

| Brain areas | Laterality | Peak T-value | Cluster size | Peak coordinate (MNI) |  |  |
| --- | --- | --- | --- | --- | --- | --- |
|  |  |  |  | x | y | z |
| Anterior Supramarginal Gyrus | L | -3.93 | 303 | -52 | -29 | 40 |
| Anterior Supramarginal Gyrus | R | -2.66 | 18 | 67 | -29 | 27 |
| Superior Frontal Gyrus | L | -4.53 | 2766 | -12 | 4 | 66 |
| Superior Frontal Gyrus | R | -4.86 | 2122 | 12 | 21 | 59 |
| Middle Frontal Gyrus | L | -5.57 | 3320 | -44 | 26 | 33 |
| Middle Frontal Gyrus | R | -5.56 | 3312 | 40 | 25 | 37 |
| Anterior Paracingulate Gyrus | L | -7.28 | 4153 | -5 | 38 | -10 |
| Anterior Paracingulate Gyrus | R | -7.13 | 3918 | 4 | 45 | -8 |
| Angular Gyrus | L | -3.52 | 519 | -62 | -54 | 26 |
| Angular Gyrus | R | -2.66 | 401 | 50 | -52 | 49 |
| Frontal Orbital Cortex | L | -6.46 | 6392 | -38 | 33 | -17 |
| Frontal Orbital Cortex | R | -7.26 | 5637 | 28 | 28 | -20 |
| Superior Parietal Lobule | L | -5.10 | 1774 | -35 | -44 | 58 |
| Superior Parietal Lobule | R | -5.96 | 1457 | 35 | -41 | 59 |
| Insular Cortex | L | -4.89 | 1004 | -32 | -26 | 16 |
| Insular Cortex | R | -6.34 | 1028 | 34 | -24 | 9 |
| Posterior Cingulate Gyrus | L | -10.83 | 4525 | -5 | -47 | 19 |
| Posterior Cingulate Gyrus | R | -10.47 | 5838 | 5 | -43 | 27 |
| Posterior Supramarginal Gyrus | L | -3.24 | 102 | -61 | -51 | 26 |
| Posterior Supramarginal Gyrus | R | -2.41 | 7 | 67 | -38 | 21 |

**Table S6.** Peak and significant cluster of fMRI activity during spindle main effect. We used the Spindle main effect whole-brain activation patterns in **Fig. 3c**. ROIs were defined anatomically (see Methods). Cluster sizes are reported *p_unc._* < 0.001.

| Brain areas | Laterality | Peak T-value | Cluster size | Peak coordinate (MNI) |  |  |
| --- | --- | --- | --- | --- | --- | --- |
|  |  |  |  | x | y | z |
| Anterior Supramarginal Gyrus | L | -6.36 | 1357 | -55 | -29 | 45 |
| Anterior Supramarginal Gyrus | R | -6.53 | 1671 | 55 | -26 | 49 |
| Superior Frontal Gyrus | L | -6.83 | 2274 | -20 | 33 | 38 |
| Superior Frontal Gyrus | R | 5.52 | 2320 | 12 | -4 | 75 |
| Middle Frontal Gyrus | L | -8.03 | 5350 | -42 | 30 | 30 |
| Middle Frontal Gyrus | R | -5.77 | 4194 | 31 | 28 | 41 |
| Anterior Paracingulate Gyrus | L | 7.15 | 1086 | -7 | 11 | 42 |
| Anterior Paracingulate Gyrus | R | 6.68 | 2222 | 2 | 15 | 41 |
| Angular Gyrus | L | -7.55 | 1240 | -48 | -58 | 50 |
| Angular Gyrus | R | -7.33 | 4588 | 53 | -53 | 34 |
| Frontal Orbital Cortex | L | -4.61 | 1149 | -41 | 21 | -11 |
| Frontal Orbital Cortex | R | -3.98 | 645 | 40 | 20 | -16 |
| Superior Parietal Lobule | L | -6.91 | 1687 | -36 | -56 | 61 |
| Superior Parietal Lobule | R | -7.79 | 3091 | 42 | -43 | 62 |
| Insular Cortex | L | -6.99 | 1103 | -37 | -20 | 15 |
| Insular Cortex | R | -7.26 | 2169 | 36 | -22 | 12 |
| Posterior Cingulate Gyrus | L | -7.85 | 3208 | -5 | -46 | 38 |
| Posterior Cingulate Gyrus | R | -7.10 | 3352 | 0 | -45 | 36 |
| Posterior Supramarginal Gyrus | L | -6.71 | 1263 | -54 | -49 | 39 |
| Posterior Supramarginal Gyrus | R | -6.61 | 2376 | 48 | -38 | 54 |

**Table S7.** Peak and significant cluster of fMRI activity during SO-spindle interaction. We used the SO-spindle interaction effect whole-brain activation patterns in **Fig. 3d**. ROIs were defined anatomically (see Methods). Cluster sizes are reported *p_unc._* < 0.001.

| Brain areas | Laterality | Peak T-value | Cluster size | Peak coordinate (MNI) |  |  |
| --- | --- | --- | --- | --- | --- | --- |
|  |  |  |  | x | y | z |
| Anterior Supramarginal Gyrus | L | -4.95 | 1327 | -56 | -33 | 38 |
| Anterior Supramarginal Gyrus | R | -2.42 | 10 | 60 | -23 | 27 |
| Superior Frontal Gyrus | L | -3.50 | 398 | -14 | 10 | 64 |
| Superior Frontal Gyrus | R | -4.44 | 596 | 6 | 23 | 49 |
| Middle Frontal Gyrus | L | -4.74 | 2455 | -50 | 11 | 40 |
| Middle Frontal Gyrus | R | -6.01 | 1694 | 53 | 17 | 37 |
| Anterior Paracingulate Gyrus | L | -3.56 | 381 | -1 | 21 | 47 |
| Anterior Paracingulate Gyrus | R | -4.98 | 725 | 7 | 21 | 47 |
| Angular Gyrus | L | 1.71 | 0 | -62 | -54 | 26 |
| Angular Gyrus | R | 2.63 | 149 | 55 | -58 | 19 |
| Frontal Orbital Cortex | L | -4.72 | 1492 | -47 | 26 | -6 |
| Frontal Orbital Cortex | R | -4.24 | 872 | 40 | 27 | -2 |
| Superior Parietal Lobule | L | -4.83 | 1177 | -37 | -46 | 56 |
| Superior Parietal Lobule | R | -4.02 | 584 | 28 | -50 | 54 |
| Insular Cortex | L | -4.01 | 1385 | -38 | 19 | -1 |
| Insular Cortex | R | -4.36 | 1374 | 32 | 21 | 0 |
| Posterior Cingulate Gyrus | L | -2.93 | 83 | -9 | -32 | 40 |
| Posterior Cingulate Gyrus | R | -2.91 | 130 | 7 | -30 | 42 |
| Posterior Supramarginal Gyrus | L | -1.96 | 0 | -48 | -44 | 46 |
| Posterior Supramarginal Gyrus | R | -2.02 | 0 | 49 | -37 | 51 |

## Notes

### Competing Interest Statement

The authors have declared no competing interest.

### Summary of Updates

This is a revised manuscript following comments and recommendations from reviewers and editors at eLife.

